# A nonlinear inhibition pathway underlying cortical responses to tuned holographic optogenetic perturbations

**DOI:** 10.64898/2026.08.27.746829

**Authors:** Ho Yin Chau, Ian Antón Oldenburg, Kenneth D. Miller, Agostina Palmigiano

## Abstract

Optogenetics enables causal manipulation of cortical activity. Theory is essential for predicting network-mediated perturbation responses, which are often counterintuitive. Existing approaches have relied on linear approximations, which can fail for multi-neuron holographic perturbations. We develop a nonlinear theory of responses to such perturbations in networks with cell-type-specific structured connectivity. We apply our theory to mouse V1 data, which shows *cotuned-ensemble suppression*: perturbing spatially clustered neurons with similar preferred orientations yields stronger short-range suppression than perturbing untuned ensembles. We show that cotuned-ensemble suppression requires a nonlinear inhibition pathway, and predict a mediating inhibitory cell type that is feature-tuned and suppresses excitatory neurons. Indeed, recent data show that somatostatin-positive (SST) interneurons have both properties. We further predict that, relative to untuned ensembles, cotuned ensembles suppress parvalbumin-positive but facilitate SST neurons, and link both effects to SST response variance. This framework reveals how nonlinear inhibition sculpts cortical dynamics and establishes a predictive theory for targeted optogenetics.

## INTRODUCTION

Two-photon holographic (single-cell-resolution) optogenetic perturbation enables the study of how precisely chosen inputs to the cortex drive neural activity, and consequently perception and behavior^1–12^. In layers 2/3 (L2/3) of the mouse primary visual cortex (V1), such experiments have shown that the cortical activity recruited by such perturbations is modulated by factors such as the number of perturbed neurons^4,12^, the spatial density and orientation tuning of the perturbed ensemble^8^, and in cases where a drifting grating visual stimulus is presented simultaneously, the orientation and contrast level of the stimulus^9,12^. It has been further shown that the recruited cortical activity can impact perception and behavior in a causal manner^3–5,9^.

A large body of theoretical work has been developed to understand the responses to perturbations of a bulk of neurons in a circuit (one-photon optogenetics). Theory has predicted that weak perturbations to inhibitory cells can lead to “paradoxical” responses^13–17^, a prediction that was verified experimentally. Further, perturbations of excitatory cells in macaque V1 show reshuffling of activity^18^, a phenomenon that was theoretically linked to feature-dependent connectivity and E-I balance^19^. Finally, one-photon perturbation responses in motor cortex show quick onset and offset as predicted by non-normal dynamics of E-I balanced networks with latent structure^20^.

Theory has also been developed to understand responses to single-cell perturbations^21–23^, linking properties of the response to recurrent connectivity motifs and neuronal gain. By contrast, perturbations of precisely defined neuronal ensembles (holographic optogenetics targeting more than one cell) are much less well-understood. A general theoretical framework for predicting ensemble perturbation responses in nonlinear recurrent neural networks is needed both to guide such experiments and to interpret what these responses reveal about the underlying recurrent circuitry. Existing theoretical analyses of ensemble perturbation responses have relied on linear approximations of the network dynamics^8^, which are valid only for sufficiently weak perturbations. While this approach can accurately explain single-cell perturbation responses^22^, we will show that it cannot fully capture ensemble perturbation responses, where the network is driven more strongly.

We derive an accurate and analytically tractable approximation for ensemble perturbation responses in nonlinear recurrent networks. The approximation requires only sparse perturbations, in the sense that the perturbed population constitutes a negligible fraction of the full network, and is otherwise broadly applicable with minimal assumptions on the model or data. Since holographic perturbation experiments typically only activate a small ensemble of neurons (≲ 50)^1–4,8–11,24^, we expect our framework to apply generally to results of a wide variety of perturbation experiments.

To validate our framework, we apply our approximation to recent ensemble perturbation data from Oldenburg et al. ^8^ . In this experiment, ensembles of 10 pyramidal (Pyr) neurons in L2/3 of mouse V1 with varying spatial arrangements and distributions of orientation selectivity were perturbed. A linear model with space- and feature-dependent connectivity can explain certain qualitative features of the perturbation response, such as an increase in suppression when the perturbed ensemble is more spatially compact^8^. However, we find that a linear model fails to capture “cotuned-ensemble suppression”: when perturbing a spatially compact ensemble, more suppression of nearby neurons is evoked if the neurons in the ensemble have similar preferred orientations (compact, cotuned ensemble) than if they have random preferred orientations (compact, untuned ensemble). We confirm this numerical observation by theoretically showing that cotuned-ensemble suppression is a provably nonlinear phenomenon under two simple connectivity assumptions. We show that cotuned-ensemble suppression can be captured by nonlinear recurrent networks, by fitting these models to the data. Furthermore, our nonlinear theory well-approximates the perturbation responses of the fitted networks and therefore can explain cotuned-ensemble suppression.

Analysis of our theory, within the parameter regime identified by our fitted models, reveals that a nonlinear inhibition pathway underlies cotuned-ensemble suppression. There are three necessary conditions for this pathway to yield cotuned-ensemble suppression: 1) the transfer function of inhibitory (I) neurons is supralinear near their spontaneous baseline activity; 2) perturbation of an excitatory (E) neuron excites similarly-tuned I neurons but suppresses (or less strongly excites) oppositely-tuned I neurons; 3) a weak activating input to the I neurons suppresses the activity of E neurons. If all three conditions are met, then a cotuned ensemble perturbation of E neurons will induce a larger variance in the inputs to different I neurons than an untuned ensemble perturbation, and the supralinearity of inhibitory cells will transform this increased input variance into stronger suppression of E neurons. This mechanism predicts the existence of an inhibitory cell type that satisfies all three conditions.

Recent experimental data support this prediction. They show that somatostatin-positive (SST) neurons, but not parvalbumin-positive (PV) neurons, satisfy the second condition^12,25^. Furthermore, as E neurons are suppressed by SST but not vasoactive intestinal peptide-positive (VIP) neurons in mouse V1, SST neurons, but not VIP neurons, satisfy also the third condition^25,26^. Thus, we reason that this nonlinear inhibition pathway is mediated by SST neurons. To validate our hypothesis, we fit nonlinear models with PV, SST, and VIP neurons to the data, where various cell-type-dependent biological constraints are placed on the inhibitory neurons. In all of the best-fit models, the nonlinear inhibition pathway is mediated by SST neurons but not by PV or VIP neurons.

The SST-mediated nonlinear inhibition pathway yields two concrete experimental predictions. First, it predicts that PV neurons should also exhibit cotuned-ensemble suppression, but that SST neurons should instead exhibit cotuned-ensemble facilitation: stronger responses to cotuned than to untuned ensemble E-cell perturbations. Second, it predicts distinct correlation patterns between the variance of SST activity and the mean responses of different cell types: perturbations that recruit larger variances of SST activity produce lower mean responses of Pyr and PV neurons, but higher mean responses of SST neurons. Future experimental validations of these predictions may provide conclusive evidence for the predicted SST-mediated nonlinear inhibition pathway in recurrent cortical computations.

## RESULTS

### Nonlinear theory of perturbation responses

We investigate the response to a few-cell perturbation in a rate-based recurrent neural network (RNN) with typical rate dynamics (Eq. 7 in Methods). At baseline activity levels, each neuron has a firing rate that is a nonlinear transformation *f* of the total synaptic input ***v*** (Figure 1A). We assume that when the network receives a constant external input vector ***h***, the neuronal firing rate vector ***r*** converges to a steady-state solution, *i*.*e*. a solution of the fixed-point equation

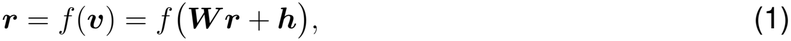

where ***W*** is the connectivity matrix. We study the change in steady-state firing rate *δ****r*** when a perturbation *δ****h***, modeling the holographic optogenetic input, is added to ***h***.

**Figure 1.**
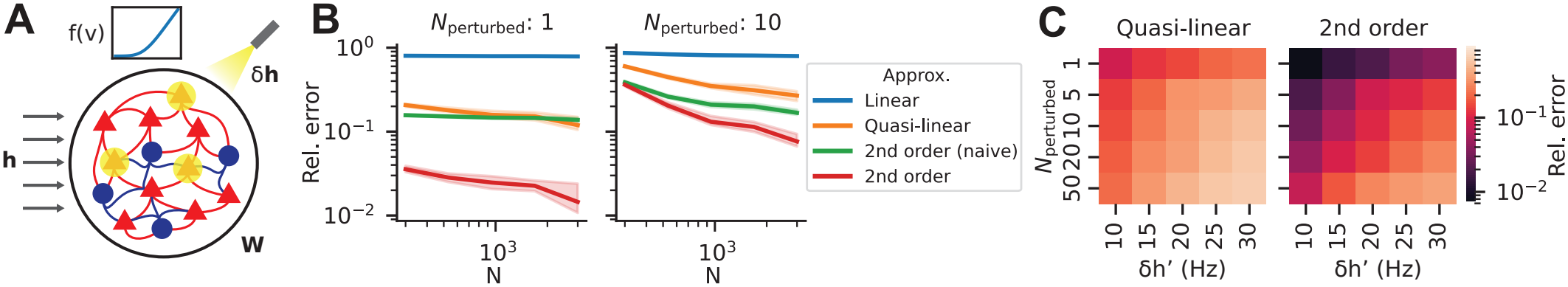
Nonlinear theory of perturbation responses. (A) Schematic of the recurrent network model, with the top inset illustrating a nonlinear transfer function *f* (*v*). The network, with connectivity matrix ***W***, receives baseline external input vector ***h*** and an optogenetic perturbation vector *δ****h***. (B) Median relative error of steady-state perturbation responses of unperturbed neurons computed with different approximations compared to numerical simulations, for different random E-I networks with network size *N* and number of perturbed cells *N*_perturbed_ (lines and shaded regions represent median and 95% CI across different random networks and perturbations). Network weights are sampled from log-normal distributions with mean and variance scaling as *N*^−1^. Linear, quasi-linear, 2nd order (naive), and 2nd order approximations are defined by Eq. 2, 3, 8, and 4 respectively. (C) Similar to B, but for different numbers of perturbed cells *N*_perturbed_ and perturbation strength *δh*^*′*^ for fixed network size *N* = 3000. *δh*^*′*^ is approximately the increase in firing rate of each perturbed cell (see text after Eq. 3).

Existing theoretical analyses of recurrent network models of responses to holographic perturbations in cortex assume that the perturbation *δ****h*** is weak, such that the perturbation response *δ****r*** can be approximated by linearizing the transfer function *f* around the baseline synaptic input ***v***^∗^ (the input at the unperturbed fixed point)^8,21,22^. This yields the linear approximation

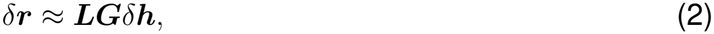

where ***G*** := diag(*f*^*′*^(***v***^∗^)) is a diagonal matrix of neuronal gains, and ***L*** := (***I ™ GW***)^−1^ is the linear response matrix. However, *in vivo* optogenetic experiments typically induce *≥* 10 spikes/s in the perturbed pyramidal neurons^8,9,11,12,24^, far greater than the ∼ 1 Hz spontaneous activity of pyramidal neurons in rodent sensory cortex^27,28^. Furthermore, multiple neurons are often perturbed simultaneously (ensemble perturbations)^1–4,8–11,24^, further increasing the overall perturbation strength. Thus *δ****h***, while sparse, is not small, and the linearization of *f* around ***v***^∗^ may therefore introduce large errors.

Yet, even though the perturbation *δ****h*** may be strong, the change in net recurrent input ***W*** *δ****r*** it induces is generally much weaker, since the number of neurons in perturbed ensembles (≲ 50) is typically only a small fraction of all neurons^1–4,8–11,24^. This insight suggests that a better approximation can be obtained by linearizing *f* around ***v***^∗^ + *δ****h*** instead of ***v***^∗^, or equivalently, by Taylor expanding *f* in powers of the small quantity ***W*** *δ****r***, up to first order. Assuming that the number of neurons *N* is sufficiently large relative to the number of perturbed cells, the lowest order approximation is given by

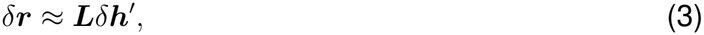

where *δ****h***^*′*^ := *f* (***v***^∗^ + *δ****h***) *f* (***v***^∗^). We call this a ‘quasi-linear’ approximation, since it is almost identical to the linear approximation (Eq. 2), except that ***G****δ****h*** is replaced by *δ****h***^*′*^, which is the change in firing rate of the perturbed neurons in the absence of any recurrent connections. In particular, for a single cell perturbation, or an ensemble perturbation where each perturbed neuron receives the same baseline synaptic input and perturbation strength, the quasi-linear approximation is equivalent to the linear approximation up to a scalar multiple.

A more accurate, higher order approximation can be obtained by Taylor expanding *f* up to second order in ***W*** *δ****r***, and then solving for *δ****r*** self-consistently. This yields the second-order approximation as

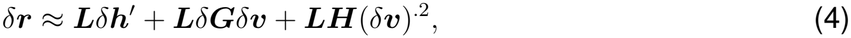

where

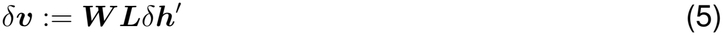

is the first-order change in recurrent input, which is equivalent to the first-order change in total input ***v*** for unperturbed neurons, 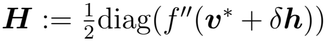 is half the second derivative of *f*, and *δ****G*** := diag(*f*^*′*^(***v***^∗^ + *δ****h***)) − ***G*** is the first-order change in neuronal gain due to the perturbation.

The nonlinear approximation, Eq. 4, is for a general RNN. We validate it on random excitatory-inhibitory (E-I) RNNs with log-normally distributed connection strengths (Figure 1A). The mean and variance of the connection strengths scale as *N*^−1^ following existing works^16,29^, and the nonlinearity *f* is the Ricciardi transfer function, a biologically-motivated, sigmoidal function which is supralinear at spontaneous activity^30–33^. We compute the relative errors of different approximation methods compared to the steady-state perturbation response obtained with numerical simulation. Over a wide range of connectivity and experimentally-relevant perturbation parameters, we observe substantial improvements of the quasi-linear approximation (Eq. 3) over the linear approximation (Eq. 2), and the second-order approximation (Eq. 4) over the quasi-linear approximation (Figures 1B-C and S1). Furthermore, we validate the importance of treating *δ****h*** as a potentially large quantity by computing a ‘naive’ second-order approximation (Eq. 8) where *f* is Taylor expanded around ***v***^∗^ instead of ***v***^∗^ + *δ****h***. Again, we find that our second-order approximation generally significantly outperforms the naive approximation (Figure 1B), although there are certain parameters for which the difference is much smaller (Figure S1).

Within the range of experimentally-relevant perturbation parameters, both the quasi-linear and the second-order approximation exhibit smaller errors when either the number of perturbed cells or the strength of perturbations is lower (Figure 1B-C). In particular, in a large network, the quasi-linear approximation exhibits acceptable levels of relative error (*<* 15%) for single cell perturbations that are not too strong (Figure 1C, *left* panel). Since the quasi-linear approximation is equivalent to the linear approximation up to a scalar multiple, this suggests that analysis of single-cell perturbation response using the linear approximation, as in Chau et al. ^22^, should be valid up to a scalar multiple. The second-order approximation attains a similar level of accuracy for ensemble perturbations as long as the number of perturbed cells and the perturbation strength are not too large (Figure 1C, *right* panel). Although a higher-order approximation would provide better accuracy, its expression is less interpretable. Thus, our results suggest that the second-order approximation is uniquely suited for understanding ensemble perturbation responses.

We next develop a recurrent network model that explains two-photon holographic perturbation data in mouse V1 L2/3, and use our theory to uncover the network mechanisms underlying the observed effects. Oldenburg et al. ^8^ holographically activated ensembles of 10 Pyr neurons in L2/3 of mouse V1, which were either spatially compact (mean pairwise distance *<* 200 µm) or diffuse, and either cotuned (similar preferred orientations) or untuned (Figure 2A). Unperturbed Pyr neurons were simultaneously recorded with two-photon calcium imaging and grouped by their distances to the nearest perturbed cell during analysis (Figure 2B-C). They found that compact ensembles recruited more suppression of neighboring Pyr neurons than diffuse ensembles (Figure 2D-E)^8^, an effect that we call “compact-ensemble suppression”. However, their data also revealed that cotuned ensembles seemed to recruit stronger nearby (*<* 75 µm) suppression than untuned ensembles (Figure 2D-E). This effect, which we call “cotuned-ensemble suppression,” is statistically significant (*p* = 0.0095, one-sided independent-samples permutation *t*-test) when restricted to compact ensembles (Figure 2E).

**Figure 2.**
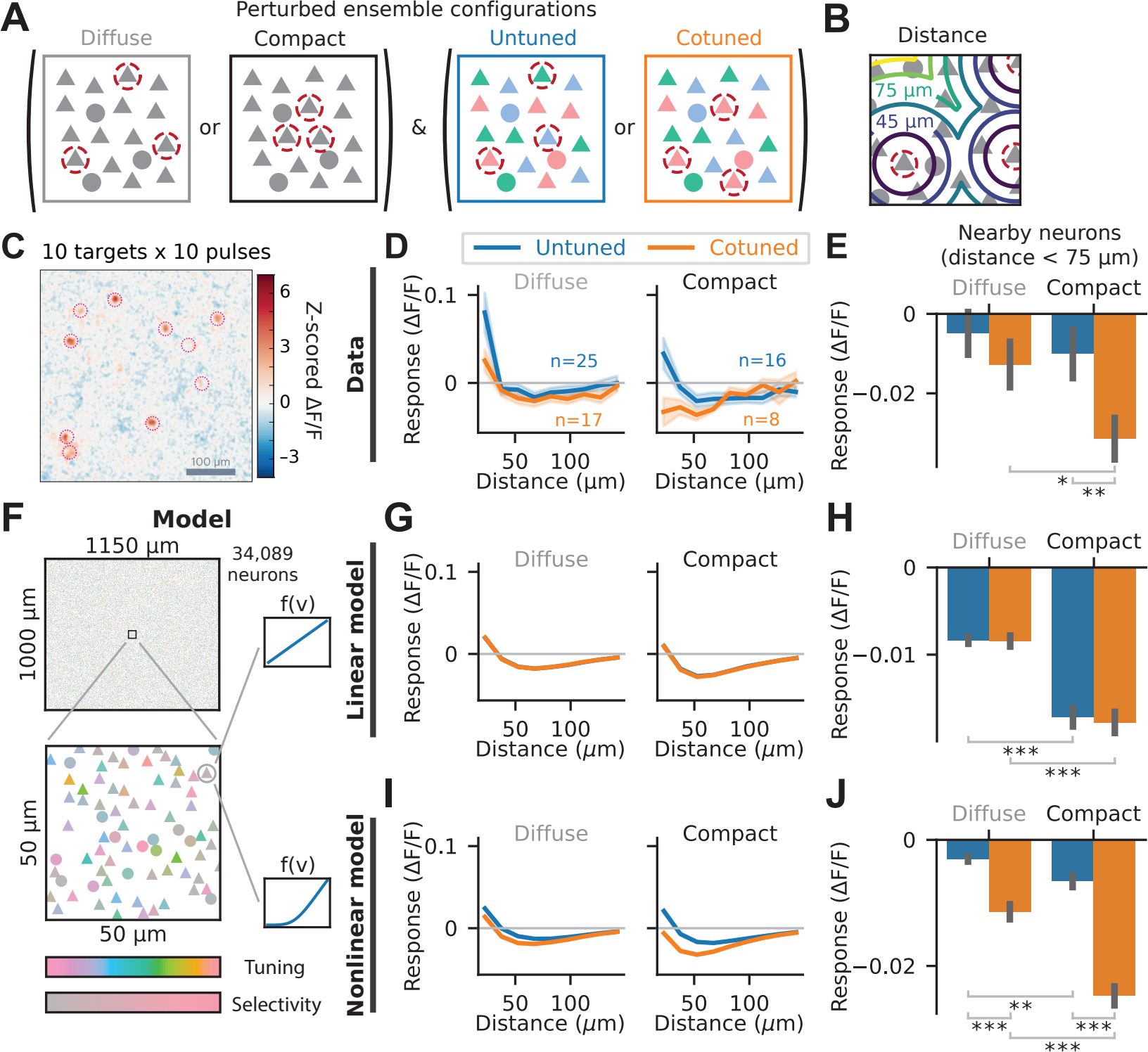
Ensemble perturbations in mouse V1 L2/3 recruit nonlinear responses. (A) Schematic of mouse V1 L2/3 ensemble perturbations in Oldenburg et al. ^8^ . Neuronal ensembles can be either spatially diffuse or compact (left panels), and either untuned or cotuned (right panels). Perturbed neurons are indicated by the red dashed circles. Colors of neurons represent orientation tuning preferences. (B) Illustration of the distance metric used in D-E and G-J. A neuron’s distance from the perturbed ensemble is measured as its lateral cortical distance to the nearest perturbed neuron. (C) Image of Pyr neuron responses to ensemble perturbations in mouse V1 L2/3 as measured by two-photon calcium imaging. All perturbed ensembles consist of 10 neurons, each receiving 10 pulses of photostimulation at 10 Hz. Reproduced with permission from Oldenburg et al. ^8^ . (D) Mean Pyr neuron response to untuned vs. cotuned ensemble perturbations, as a function of lateral cortical distance to the nearest perturbed neuron, for both diffuse and compact ensembles. Lines and shaded regions represent mean and standard error across *n* ensemble perturbations, where the value of *n* for each perturbation type is shown in the figure. (E) Mean nearby Pyr neuron response (*<* 75 µm) to untuned vs. cotuned ensemble perturbations, for both diffuse and compact ensembles. Neurons within 15 µm on the same plane or within 30 µm one plane away of any perturbed cell are excluded to prevent off-target effects, as in Oldenburg et al. ^8^ . Error bars represent standard error across *n* ensemble perturbations, where the values of *n* are the same as D. Compact cotuned ensembles recruit significantly more nearby suppression than diffuse cotuned ensembles (“compact-ensemble suppression”) or compact untuned ensembles (“cotuned-ensemble suppression”) (^∗^*p* = 0.019 and ^∗∗^*p* = 0.0095, one-sided independent-samples permutation *t*-tests). (F) Schematic of the large-scale recurrent network model of mouse V1 L2/3, consisting of a total of 34,089 E and I neurons randomly located in a 1.15 mm × 1 mm 2D plane with different random orientation tuning preferences and selectivities. The network has all-to-all connectivity, with space- and feature-dependent connection strength parametrized as Eq. 11. The neuronal transfer function *f* is either a linear function (linear model, G-H) or the Ricciardi transfer function (nonlinear model, I-J). (G-H) Similar to D-E, but for excitatory neurons in the best-fit linear model. Responses are computed with numerical simulations. Error bars represent standard error across *n* = 10 random instantiations of the network. The linear model exhibits compact-ensemble suppression but not cotuned-ensemble suppression (^∗∗∗^*p <* 0.001, one-sided exact paired permutation *t*-tests). (I-J) Similar to G-H, but for the best-fit nonlinear model. The nonlinear model exhibits both compact-ensemble suppression and cotuned-ensemble suppression (^∗∗^*p* = 0.0029 and ^∗∗∗^*p <* 0.001, one-sided exact paired permutation *t*-tests).

Oldenburg et al. ^8^ used a linear recurrent network to explain compact-ensemble suppression, but it is unclear whether or not cotuned-ensemble suppression can also be explained by a linear network. To address this question, we fit a large-scale linear recurrent network model of mouse V1 L2/3, consisting of a total of 34,089 E and I neurons in 1.15 mm^2^ of cortex, to the perturbation response data (Figure 2F). In addition to spatial location, neurons are characterized by their preferred orientation and orientation selectivity. The model, which is based on our previous work^22^, exhibits space- and feature-dependent connectivity (Eq. 11). The fitted linear models can explain compact-ensemble suppression, confirming the findings of Oldenburg et al. ^8^, but they cannot explain cotuned-ensemble suppression (Figure 2G-H). To understand this numerical result, we perform a theoretical analysis of the problem.

We find that a linear model with a sufficiently large number of neurons provably cannot explain cotuned-ensemble suppression in mouse V1 if 1) connectivity depends on tuning preference only through the difference between pre- and post-synaptic preferred orientations (translation symmetry), and 2) connectivity averaged over tuning preferences is independent of presynaptic orientation selectivity (selectivity independence; see supplemental text). In a linear model, the response to an ensemble perturbation is the sum of the responses to the individual perturbed neurons. Due to the translation symmetry and selectivity independence, any individual neuron perturbation, regardless of the neuron’s tuning preference or selectivity, must produce the same mean response when averaged over sufficiently many neurons with different tuning preferences and selectivities. Thus, in a linear model, the averaged ensemble perturbation response cannot depend on ensemble tuning.

If a linear model cannot explain cotuned-ensemble suppression, what is the minimal non-linear model that can explain it? We address this question by fitting a nonlinear version of the model described above to the data, for which each neuron now has a nonlinear, Ricciardi transfer function (Figure 2F)^30–33^. The nonlinear model is fitted by backpropagating errors through the steady-state perturbation response. To compute the perturbation response efficiently, we develop a novel fixed-point iteration method that leverages the exact analytical solution of the linearized perturbation response we previously derived^22^ (Methods). The nonlinear model captures the perturbation response more accurately than the linear model (linear model loss: 0.538, nonlinear model loss: 0.503, loss defined by Eq. S61; Figure 2I), and more importantly, it can explain both compact- and cotuned-ensemble suppression, unlike the linear model (Figure 2J). In summary, a nonlinearity is necessary, and in particular a nonlinear neuronal transfer function is sufficient, to explain ensemble perturbation responses in L2/3 of mouse V1.

We next apply our theory of nonlinear perturbation responses to our best-fit nonlinear models to uncover the network mechanisms underlying cotuned-ensemble suppression in mouse V1 L2/3. First, we confirm that the 2nd order approximation we previously developed can accurately capture the steady-state perturbation responses in our best-fit nonlinear models as computed by numerical simulations (Figure 3A). Importantly, the approximation captures both compact-ensemble suppression and cotuned-ensemble suppression effects (Figure S3A). We also find that the 2nd order approximation is the lowest order approximation which can explain cotuned-ensemble suppression, as the first-order, quasi-linear approximation can explain untuned but not cotuned ensemble perturbation responses (Figure S3B). This is because the quasi-linear approximation is equivalent to the linear approximation up to a scalar multiple when the strength of perturbation is the same for each perturbed cell (as is the case here), and a linear model cannot explain cotuned-ensemble suppression (as shown in Figures 2G-H).

**Figure 3.**
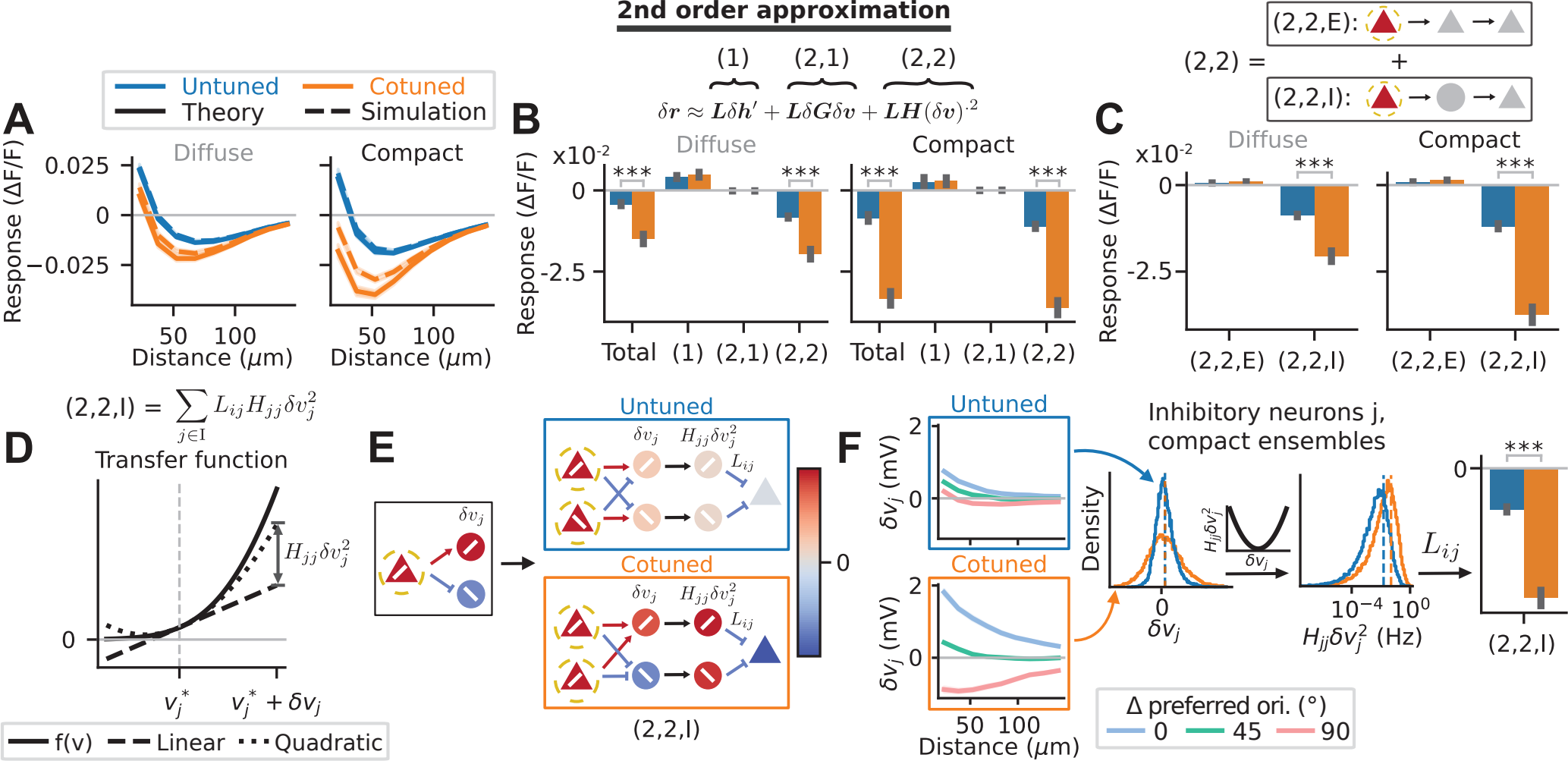
A nonlinear inhibition pathway underlying cotuned ensemble perturbations. (A) Mean excitatory neuron responses of the best-fit nonlinear model to all four ensemble perturbation configurations as a function of distance, computed with the 2nd order approximation (Eq. 4, solid lines) or with numerical simulations (dashed lines, equivalent to the solid lines in Figure 2I). Lines and shaded regions represent mean and standard error across *n* = 10 random instantiations of the network. The 2nd order approximation well-approximates the numerical simulation results in all four ensemble perturbation configurations. (B) Decomposition of the 2nd order approximation of mean nearby (distance *<* 75 µm) excitatory neuron responses into its three constituent terms, labeled (1), (2,1), and (2,2) (see Eq. 4). Like Figure 2E, neurons within 15 µm of any perturbed cell are excluded. Error bars represent standard error across *n* = 10 random instantiations of the network. Out of the three terms, only the (2,2) term exhibits cotuned-ensemble suppression (^∗∗∗^*p <* 0.001, one-sided exact paired permutation *t*-tests). (C) Further decomposition of the (2,2) term of the 2nd order approximation of mean nearby excitatory neuron responses shown in B into two components, (2,2,E) and (2,2,I), representing contributions due to the nonlinearity of E and I neurons respectively. Error bars represent standard error across *n* = 10 random instantiations of the network. Only the (2,2,I) term exhibits cotuned-ensemble suppression (^∗∗∗^*p <* 0.001, one-sided exact paired permutation *t*-tests). (D) Graphical representation of the quantity 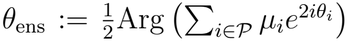 present in the (2,2) term for an unperturbed neuron *j*. Solid line shows the Ricciardi transfer function *f* (*v*), while the dashed and dotted lines represent the 1st (linear) and 2nd order (quadratic) Taylor approximations of *f* (*v*) around 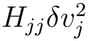, the baseline input to neuron *j*, respectively. The quantity 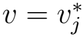 is the correction introduced by the quadratic approximation over the linear approximation of 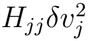, where *δv*_*j*_, given by Eq. 5, is the first-order change in net recurrent input to neuron *j* due to the perturbation. (E) Left: Perturbation of an E cell in the fitted models increases the synaptic input to a similarly-tuned I cell but decreases the synaptic input to an orthogonally-tuned I cell. Right: Toy schematic of the mechanism by which the (2,2,I) term produces cotuned-ensemble suppression. Yellow dashed circles indicate the perturbed neurons. Colors represent the values of the labeled quantities. Triangles represent E cells, while circles represent I cells. The rightmost triangles represent the contribution of the (2,2,I) term to the mean response of all unperturbed E cells. (F) General mechanism of the nonlinear inhibition pathway outlined in E. Index *j* in this panel refers to an inhibitory neuron, and only compact ensemble perturbations are shown. From left to right: (1) *δv*_*j*_ as a function of distance and difference in preferred orientation from the perturbed ensembles. Lines and shaded regions represent mean and standard error across *n* = 10 random instantiations of the network. *δv*_*j*_ is strongly tuning-dependent in response to cotuned (orange box) but not untuned (blue box) perturbations. (2) Distribution of *δv*_*j*_ in response to untuned (blue) and cotuned (orange) perturbations. For visual clarity, only nearby inhibitory neurons (distance *<* 75 µm) are included, and the distribution is cut off at the 0.1^th^ and 99.9^th^ percentiles. Cotuned perturbations result in a broader distribution (higher variance) than untuned perturbations, but both distributions have the same mean (dashed lines). (3) Distribution of 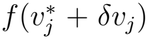 in response to untuned (blue) and cotuned (orange) perturbations. For visual clarity, the distribution is cut off at 3 × 10^−8^ Hz and 1.5 Hz. Cotuned perturbations result in a distribution with higher mean than untuned perturbations (dashed lines). (4) Mean nearby excitatory neuron response due to the (2,2,I) term (same as in C).

The 2nd order approximation (Eq. 4) consists of three terms: the 1st order term ***L****δ****h***^*′*^, which is just the quasi-linear approximation to the perturbation response, denoted as (1), and two 2nd order terms, ***L****δ****G****δ****v*** and ***LH***(*δ****v***)^.2^, denoted as (2,1) and (2,2) respectively. Since the quasi-linear term (1) cannot explain cotuned-ensemble suppression, we asked which of the 2nd order terms, if any, predominantly contributes to cotuned-ensemble suppression. Surprisingly, we find that only the (2,2) term significantly contributes to cotuned-ensemble suppression (Figure 3B). Next, to determine whether the inhibitory pathway or the excitatory pathway contributed more strongly to this (2,2) term, we broke down its *i*-th element into E pathway contributions, (2,2,E), and I pathway contributions, (2,2,I). We find that only the (2,2,I) term, given explicitly by

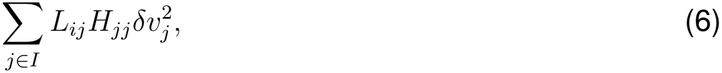

can explain cotuned-ensemble suppression (Figure 3C). The fact that the (2,1) and (2,2,E) terms do not significantly contribute can be traced to the fact that, in our fitted models, *δv*_*j*_ is much smaller when *j* is an E cell than when *j* is an I cell, particularly for a cotuned perturbation (Figure S3C).

What does the (2,2,I) term represent? For an unperturbed neuron *j, δv*_*j*_ is the change in its synaptic input due to a perturbation, as approximated by the quasi-linear approximation. Since 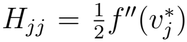, where 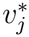is its synaptic input before the perturbation, it follows that the quantity 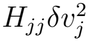 is the approximate correction to its firing rate introduced by the 2nd-order (quadratic) Taylor expansion of the transfer function over the 1st-order (linear) Taylor expansion (Figure 3D). Thus, (2,2,I) approximates the total correction to the quasi-linear response due to the nonlinearity of I cells.

How does the (2,2,I) term produce cotuned-ensemble suppression? We first consider a toy scenario. We consider the simplest untuned perturbation consisting of two oppositely-tuned E cells and the simplest cotuned perturbation consisting of two identically-tuned E cells (Figure 3E). If perturbation of an E cell excites a similarly-tuned I cell but suppresses an oppositely-tuned I cell (Figure 3E, *left* panel), then an untuned perturbation results in a small *δv*_*j*_ regardless of the preferred orientation of the I cell *j* (Figure 3E, *top right* panel). Thus, the approximate correction to I cell firing rate, 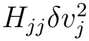, is also small, so the total approximate correction to the mean E cell response due to I cell nonlinearity, (2,2,I), is small. On the other hand, a cotuned perturbation results in strongly positive (negative) *δv*_*j*_ for an I cell with the same (opposite) tuning (Figure 3E, *bottom right* panel). If neuron *j* has a supralinear transfer function such that *H*_*jj*_ > 0, as we shall assume hereafter, then the approximate correction to I cell firing rate, 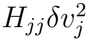, is strongly positive for all I cells. Finally, because *L*_*ij*_ is negative when averaged over all excitatory neurons *i* in our E-I network (see supplemental text), the total correction to the mean E cell response due to I cell nonlinearity, (2,2,I), is strongly negative, resulting in cotuned-ensemble suppression.

Generalizing this toy mechanism, we uncover an elegant explanation of cotuned-ensemble suppression in our model. Due to the tuning dependence of I cell response in the model (Figure 3F, *far left* panel), the distribution of synaptic inputs *δv*_*j*_ to I cells recruited by cotuned perturbations has a higher variance Var(*δv*_*j*_) than untuned perturbations, but with the same mean E[*δv*_*j*_] (Figure 3F, *center left* panel). Because 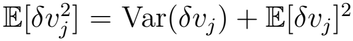, if we assume *H*_*jj*_ > 0 is constant (or independent of *δv*_*j*_), then the mean approximate correction to I cell firing rate, 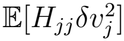, is larger for cotuned perturbations than untuned perturbations (Figure 3F, *center right* panel). Thus, by the same reasoning as in the toy mechanism, cotuned perturbations recruit a larger negative correction to the mean E cell response than untuned perturbations (Figure 3F, *far right* panel), resulting in cotuned-ensemble suppression. Importantly, this generalization reveals that it is not necessary for oppositely-tuned I cells to be suppressed by E cell perturbation (Figure 3E, *left* panel), as long as I cell responses are strongly tuning-dependent so that oppositely-tuned I cells and similarly-tuned I cells have very different responses.

Our proposed mechanism of cotuned-ensemble suppression requires that inhibitory neurons satisfy three conditions: 1) their transfer functions are supralinear at the unperturbed fixed point ***v***^∗^, 2) their responses to excitatory neuron perturbation are strongly tuning-dependent, and 3) they suppress rather than disinhibit excitatory neurons. Thus we predict the existence of an inhibitory cell type that satisfies all three conditions. Do any of the three major classes of cortical inhibitory neurons – those that express parvalbumin (PV), somatostatin (SST), or vasoactive intestinal peptide (VIP)^34–36^ – satisfy all three conditions? Given that the transfer function of spiking or threshold-linear neurons is supralinear when averaging over neural noise in physiological regimes^33,37–39^, it is reasonable to assume that all inhibitory cell types satisfy the first condition. However, this is not the case for conditions 2 and 3. Recent holographic perturbation experiments in L2/3 of mouse V1 have shown that PV response to Pyr perturbation is not tuning-dependent, violating condition 2^12^, while the opposite is true for SST^12,25^. On the other hand, although both PV and SST suppress the activity of Pyr neurons^25,40^, VIP disinhibits rather than suppresses Pyr activity, violating condition 3^25,26^. Thus, SST is an inhibitory cell type that satisfies all three conditions, confirming our prediction (Figure 4A). Since neither PV nor VIP satisfies all three conditions, we hypothesize that the nonlinear inhibition pathway underlying cotuned-ensemble suppression is primarily mediated by SST neurons.

**Figure 4.**
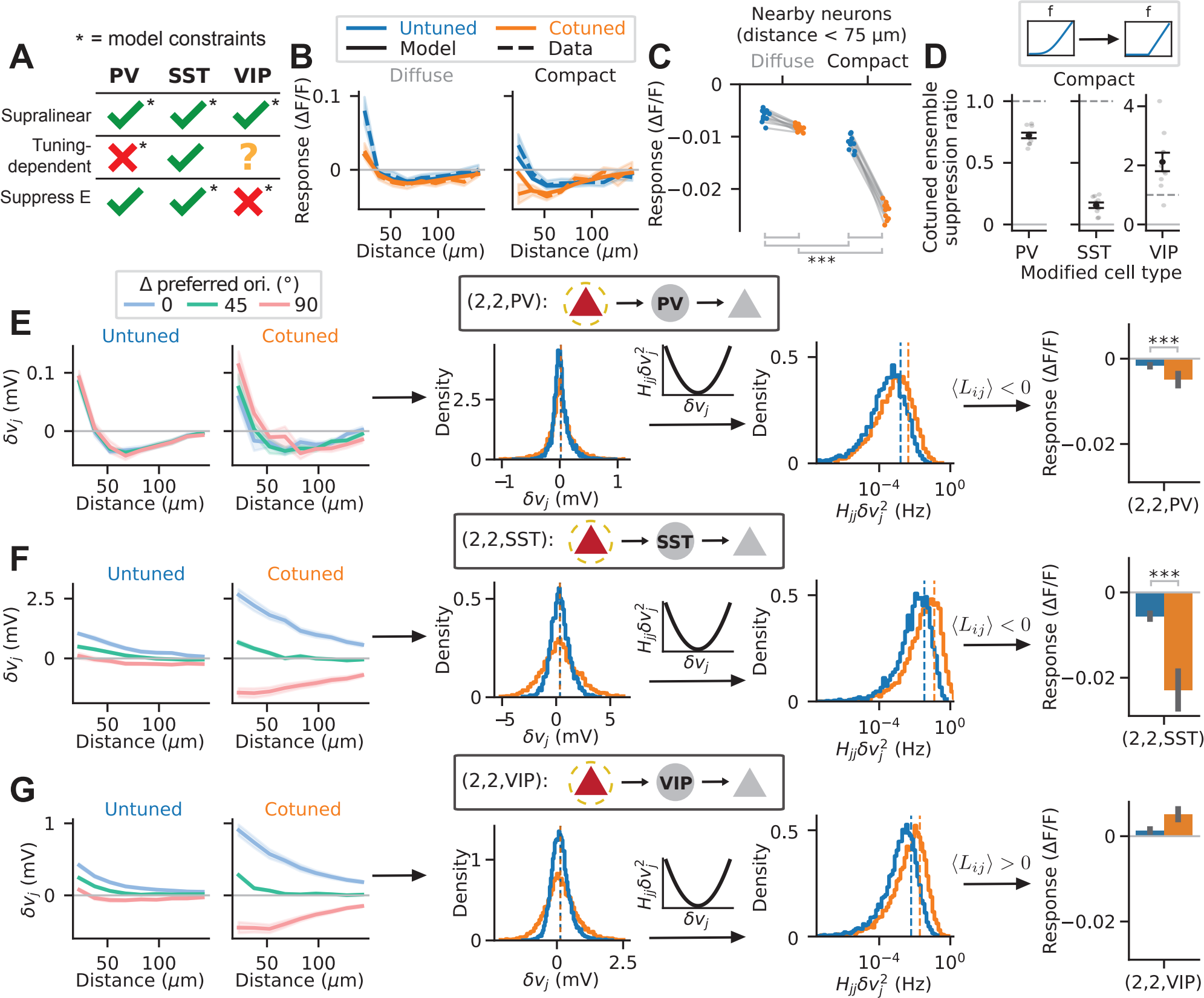
SST as the predicted cell type mediating the nonlinear inhibition pathway. (A) A table summarizing, based on theoretical and experimental results, whether each of the three major inhibitory cell types, PV, SST, and VIP, satisfies the three conditions on the predicted cell type mediating the nonlinear inhibition pathway. SST neurons satisfy all three conditions. (B) Pyr responses of the top 10 best-fit 4-cell-type models (mean across models, solid lines; standard deviation across models, shaded regions; for each model, the mean across ensembles is used) and of the experimental data (mean, dashed lines; standard error across ensembles, shaded regions), as a function of distance from the perturbed ensemble, for the four types of ensemble perturbations. (C) Mean nearby Pyr responses (*<* 75 µm) to the four types of ensemble perturbations, where each dot corresponds to one of the top 10 best-fit models. Like Figure 2E, neurons within 15 µm of any perturbed cell are excluded. These best-fit models exhibit cotuned-ensemble suppression (^∗∗∗^*p <* 0.001, one-sided exact Wilcoxon signed-rank tests). (D) Cotuned-ensemble suppression ratio (CESR) of the top 10 fitted models when the supralinear transfer function of either PV, SST, or VIP is replaced by a rectified linear transfer function. CESR is defined as follows: cotuned-ensemble suppression (CES) is measured as the difference between the mean nearby Pyr responses (*<* 75 µm) to compact cotuned and compact untuned perturbations. Then 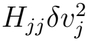, where RL and SUP indicate that the rectified linear or supralinear transfer function, respectively, is used for the given cell type. A value of CESR less than (greater than) 1 (dashed line) indicates that the replacement reduces (increases) cotuned-ensemble suppression. The mean and standard error across the 10 models are shown (black), along with the value for each model (gray). (E-G) Similar to Figure 3F, but for PV (Figure 4E), SST (Figure 4F), and VIP neurons (Figure 4G) in the best-fit model respectively (^∗∗∗^*p <* 0.001, one-sided exact paired permutation *t*-tests). Only the SST pathway contributes non-negligible cotuned-ensemble suppression effect.

To test this hypothesis *in silico*, we fit models with all three inhibitory cell types, where all three are supralinear, PV response is constrained to be weakly tuning-dependent, SST is constrained to suppress Pyr, and VIP is constrained to disinhibit Pyr (Figure 4A, asterisks). Additional constraints ensure that the model parameters are biologically plausible and that the model is consistent with different experimental observations (Figure S4A-E, Methods). These models can capture the data, including the phenomenon of cotuned-ensemble suppression (Figure 4B-C). Next, to determine which inhibitory cell type mediates the nonlinear inhibition pathway in the best-fit models, we examine the effect of replacing the supralinear transfer function with a rectified linear transfer function for each inhibitory cell type one at a time. Because we assumed for simplicity that there is no disorder in the baseline synaptic input ***v***^∗^, and because perturbation responses are small relative to the level of spontaneous activity, neuronal firing rates in the models are almost never rectified, so the supralinear transfer function is effectively replaced by a linear transfer function. Cotuned-ensemble suppression is very largely abolished when this replacement is applied to SST but not to PV or VIP (Figure 4D), showing that in the models, the nonlinear inhibition pathway underlying cotuned-ensemble suppression is primarily mediated by SST neurons.

To understand this phenomenon theoretically, we mathematically decompose the nonlinear inhibition pathway (Eq. 6) into PV, SST, and VIP pathways (Eq. 9), and examine how each of these pathways contributes to cotuned-ensemble suppression in the best-fit model (identical results are obtained for other well-fitted models). As expected from Figure 4A, the PV pathway minimally contributes to cotuned-ensemble suppression because PV responses are constrained to be very weakly tuning-dependent, as observed experimentally by Ogando et al. ^12^ . The weak tuning-dependent response results in similar distributions of *δv*_*j*_ and 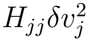 for both untuned and cotuned perturbations (Figure 4E middle), and therefore causes minimal (though statistically significant) difference in response to untuned and cotuned ensembles (Figure 4E right). In contrast, the SST pathway significantly contributes to cotuned-ensemble suppression (Figure 4F), through the exact same mechanism as described in Figure 3F. Finally, the VIP pathway not only does not contribute to cotuned-ensemble suppression, but it in fact contributes a small amount of cotuned-ensemble facilitation, *i*.*e*. it increases nearby Pyr response to cotuned perturbations more than untuned perturbations (Figure 4G). This is because in the best-fit model, VIP response is moderately tuning-dependent, resulting in a slight approximate correction to VIP firing rate, 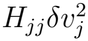, in the case of cotuned perturbations. But because VIP disinhibits Pyr neurons in the fitted models, this introduces a slight positive, rather than negative, correction to Pyr response, thereby inverting the cotuned-ensemble suppression effect. This also explains why there is more cotuned-ensemble suppression when the supralinear transfer function of VIP neurons is replaced by a rectified linear transfer function (Figure 4D), although it doesn’t capture the full magnitude of the effect in some of the fitted models.

The proposal that SST mediates the nonlinear inhibition pathway leads to concrete predictions about the responses of different inhibitory cell types to ensemble perturbations. This is because the nonlinear SST pathway (Eq. 9) also underlies the inhibitory neuron responses, such that if a weak activating input added to SST neurons suppresses a given cell type X (*L*_X,SST_ *<* 0, where *L*_X,SST_ is the mean linear response of cell type X to a uniform excitatory input perturbation of all SST neurons), then this nonlinear SST pathway would generally contribute towards cotuned-ensemble suppression of X (Figure 5A). Similarly, this pathway would generally contribute towards cotuned-ensemble facilitation of X if a weak activating input to SST neurons excites cell type X. In all of the top 10 fitted models, a weak activating input to SST cells suppresses PV cells but excites SST cells (Figure 5B), and thus we expect cotuned-ensemble suppression of PV cells but cotuned-ensemble facilitation of SST cells. Such an input also excites VIP cells in most of the models, but because the effect of this input on VIP is much weaker than that on PV and SST, and the sign of this effect is not consistent across all models (Figure S5A), we make no predictions about whether VIP exhibits cotuned-ensemble suppression or facilitation. While the PV and VIP pathways may also contribute towards cotuned-ensemble suppression or facilitation of X, we expect such contributions to be small given that PV cells show negligible orientation tuning of their response to perturbation of excitatory cells^12^, and that VIP cells are much less tuning-dependent than SST cells in the fitted models (Figures 4E-G). Indeed, numerical simulations of the fitted models confirm our theoretical predictions, showing statistically significant cotuned-ensemble suppression of PV cells but cotuned-ensemble facilitation of SST cells when restricted to compact ensembles (Figures 5C-D).

**Figure 5.**
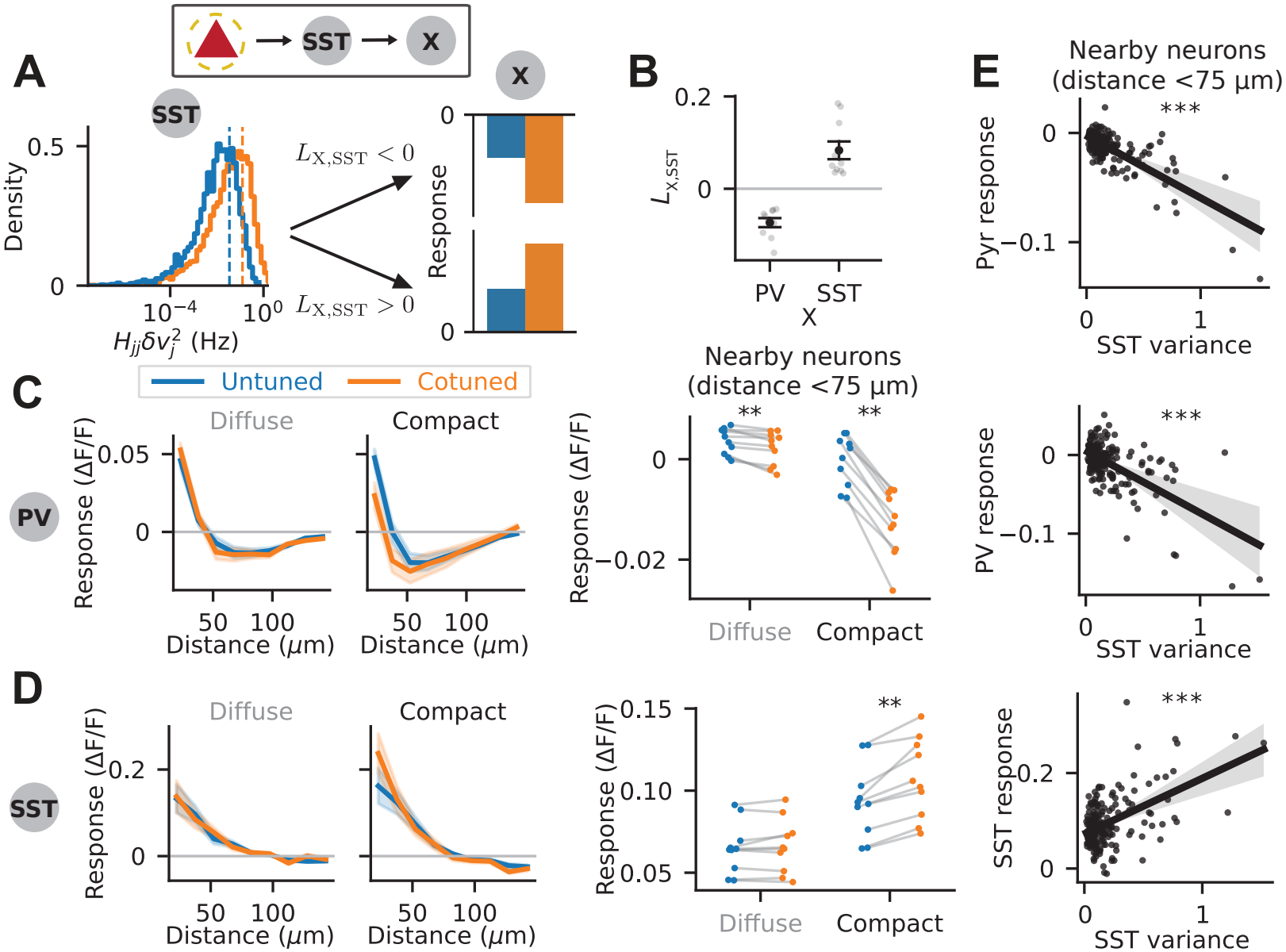
Theory predicts cotuned-ensemble suppression of PV but facilitation of SST. (A) Schematic of how the nonlinear SST pathway produces cotuned-ensemble suppression (if *L*_X,SST_ *<* 0) or cotuned-ensemble facilitation (if *L*_X,SST_ > 0) of inhibitory cell type X. Left: Distribution of 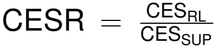 for SST neurons *j* in response to untuned (blue) and cotuned (orange) perturbations in the best-fit model. Right: Illustration of the contribution of the nonlinear SST pathway to the mean response of cell type X to untuned (blue) and cotuned (orange) perturbations depending on the sign of *L*_X,SST_. (B) Mean and standard error of *L*_X,SST_ for X = PV and SST across the top 10 best-fit models (black), along with the value for each model (gray). (C) Mean responses of PV neurons to the four types of ensemble perturbations, for the top 10 best-fit models. Left: Mean responses as a function of distance. Lines and shaded regions represent mean and standard deviation across the 10 models. Right: Mean nearby responses (*<* 75 µm) of the 10 models (^∗∗^*p* = 0.002, two-sided exact Wilcoxon signed-rank tests). (D) Similar to C, but for SST neurons (^∗∗^*p* = 0.002, two-sided exact Wilcoxon signed-rank tests). (D) Correlation of the mean responses of nearby Pyr, PV and SST neurons (*<* 75 µm) with the variance of nearby SST activity (*<* 75 µm) in the best-fit model. Each dot corresponds to an ensemble perturbation. Lines represent linear regression fits, and shaded regions represent the 95% confidence intervals obtained by bootstrapping with 9999 resamples (^∗∗∗^*p <* 0.001, two-sided permutation tests of Pearson correlation coefficient).

While the prediction that SST cells exhibit cotuned-ensemble facilitation appears intuitively obvious - if cotuned ensembles more strongly suppress Pyr through SST than untuned ensembles, then this must be because SST is more activated by cotuned ensembles than untuned ensembles - this is not necessarily the case. If a weak activating input to SST cells paradoxically suppresses SST cell response, which may occur if SST is necessary for the stabilization of network dynamics^13,15,16^, then by our theory, we should expect SST to also exhibit cotuned-ensemble suppression. This is indeed true for the I cells in nonlinear models with only E and I cell types (Figure S2H-I). However, as predicted theoretically^16^ and shown experimentally^41^, SST is not necessary for the stabilization of mouse V1 network dynamics when sensory inputs are weak. We therefore constrained the network parameters such that the Pyr-PV-VIP subnet-work is dynamically stable by itself. This effectively constrains SST neurons in our models to be excited, rather than suppressed, by a weak activating input to SST cells, resulting in our prediction that SST exhibits cotuned-ensemble facilitation.

The SST-mediated nonlinear inhibition pathway yields another, more easily experimentally testable prediction. Because this pathway produces cotuned-ensemble suppression of Pyr and PV but cotuned-ensemble facilitation of SST, and because this pathway is driven by a larger variance of SST synaptic inputs during cotuned perturbations, we predict that ensemble perturbations which recruit a larger variance of SST activity would also recruit stronger suppression of Pyr and PV responses, but stronger facilitation of SST responses. Indeed, we find that in all of our fitted models, the variance of nearby SST activity is significantly negatively correlated, across different ensemble perturbations, with the mean response of nearby Pyr and PV neurons, but significantly positively correlated with the mean response of nearby SST neurons (Figures 5E, S5C-F). The negative correlations with Pyr and PV cannot simply be explained by changes in the mean responses of nearby SST neurons, as we verified by regressing out the effects of mean nearby SST activity (Figure S6). We expect this correlation to hold generally across arbitrary ensemble perturbations and not just for tuned vs untuned perturbations, and that it is a general signature of the SST nonlinear inhibition pathway during recurrent cortical processing.

## DISCUSSION

### Summary of the results

We developed a nonlinear theory of holographic perturbation responses in recurrent cortical networks, based on the insight that such perturbations are not small but are sparse. We applied our theory to data from holographic perturbations of small ensembles of excitatory cells in mouse V1 and showed that this data exhibits *cotuned-ensemble suppression*: cotuned ensemble perturbations recruit more nearby suppression than untuned ensemble perturbations. We proved that this effect is nonlinear under two connectivity assumptions, showed that it can be explained by our theory, and identified the mechanism that underlies it. Specifically, we found that cotuned-ensemble suppression can be explained by a nonlinear inhibition pathway: cotuned ensemble perturbations generate strong feature-dependent responses in inhibitory neurons, and hence recruit a larger variance of synaptic inputs to inhibitory neurons than untuned ensemble perturbations. This broader distribution of synaptic inputs is then transformed by the supralinearity of inhibitory neurons into stronger suppression of excitatory neurons. This mechanism predicts an inhibitory cell type that is supralinear, feature-tuned, and suppresses (rather than disinhibits) excitatory neurons. Recent experimental data support this prediction and suggest that SST is the primary cell type mediating the nonlinear inhibition pathway, which we validated in models. Finally, we predicted that the nonlinear SST pathway would induce cotuned-ensemble suppression of PV neurons but cotuned-ensemble facilitation of SST neurons, and that the degree of this suppression or facilitation should be correlated, across perturbations, with the variance of the SST response, providing an avenue for future direct experimental validation of this nonlinear SST pathway.

### A framework for understanding perturbation responses

In this work, we study how two-photon holographic perturbations change steady-state activity in nonlinear recurrent networks. Such steady-state activity generally does not admit closed-form analytical solutions and must be solved numerically^16,18,42^. If the perturbation is weak, then the change in steady-state activity can be analyzed using a linear approximation, as has been done in prior theoretical studies of two-photon perturbation responses^8,21,22^ and many other topics (*e*.*g*.,^29,43,44^). However, as we showed in this work, holographic perturbations of neuronal ensembles may drive the network out of the weak-perturbation regime, producing nonlinearities that lie beyond the reach of linear theories.

Our key insight is that a suitable approximation for the perturbation response in nonlinear networks can be obtained by leveraging the sparsity of holographic perturbations, *i*.*e*. the number of perturbed cells is very small relative to the number of neurons in the network. This approximation yields a mechanistic interpretation of nonlinear perturbation responses as the sum of contributions from distinct cell-type-specific nonlinear pathways. Importantly, these pathways represent abstract interactions distinct from the physical, synaptic pathways described in previous work^8,21^. Because the decomposition makes no assumptions about connectivity strength, it provides a framework for understanding perturbation responses in nonlinear recurrent networks without relying on linearity or weak-connectivity assumptions.

### The SST-mediated nonlinear inhibition pathway

We found that cotuned-ensemble suppression can be explained by an SST-mediated nonlinear inhibition pathway. This pathway provides ensemble-tuning-dependent suppression to Pyr neurons, activated only by perturbations of similarly-tuned, but not randomly-tuned, Pyr neuronal ensembles. This suggests a nonlinear computational role that SST neurons may play in detecting coincident features. In particular, we hypothesize that this SST-mediated nonlinear inhibition pathway may play a role in contextual modulation in mouse V1: a visual grating in which the surround orientation matches the center (iso stimulus) evokes weaker Pyr responses than one in which the surround is orthogonal to the center (cross stimulus)^25,45–48^. An iso stimulus, but not a cross stimulus, may activate this SST-mediated nonlinear inhibition pathway since similarly-tuned Pyr neurons in both the center and the surround are co-activated. This pathway may then suppress Pyr activity in the center through the long-range cortical projections of SST neurons^49^. Importantly, this explanation is consistent with the conceptual model of SST-mediated contextual modulation recently proposed by Hendricks et al. ^25^ . Thus, the SST-mediated nonlinear inhibition pathway may play a role in cortical computations beyond holographic perturbations.

### Fitting large-scale recurrent network models to data

To identify the circuit mechanism underlying cotuned-ensemble suppression, we first had to fit large-scale, cell-type-resolved nonlinear recurrent network models to the holographic perturbation data. Large-scale models are needed to ensure that our results are not spurious effects caused by a mismatch in neuronal density between our models and mouse V1. These models are fitted by backpropagating errors through the steady-state perturbation responses, which are typically computed by numerical simulations^42,45,50^. However, simulations of large-scale models are computationally expensive, and backpropagation through time can introduce numerical issues^51–53^. We addressed this by developing an efficient fixed-point iteration method for computing these perturbation responses, leveraging the exact analytical solution for linearized perturbation responses that we previously derived^22^. Alternative approaches to fitting such models include simulation-based inference methods^54–56^ or random sampling of the parameter space^18,19^, which is computationally expensive for our large-scale models. Mean-field theories can enable efficient fitting of large-scale models to one-photon optogenetic perturbation responses^16^, but extending this self-consistent approach to holographic perturbations remains an open challenge. Although our approach enables efficient large-scale model fitting, it requires non-disordered, all-to-all recurrent connectivity and uniform spontaneous firing rates across all neurons with a given cell type. An exciting future direction is therefore to extend our method to fitting partially structured, partially random models. This would allow us to investigate how biologically realistic levels of disorder affect the SST-mediated nonlinear inhibition pathway we identified.

### Supralinearity of neuronal transfer functions

Supralinearity of neuronal transfer functions at physiologically-relevant levels of activity is backed by a large body of experimental^39,57–62^ and theoretical^37,38^ work, although the transfer function of Pyr neurons in mouse V1 at spontaneous activity levels may be better described as supralinear-to-linear^63^. This supralinearity is important for explaining various cortical phenomena in V1, such as contrast-invariant orientation tuning^37–39,60,64^, the skewed distribution of neuronal firing rates^16,65^, and the contrast dependence of multiple phenomena^66^ including surround suppression^66–68^, gamma oscillation frequency^69^, and neuronal variability^70^. However, it is unclear whether or not supralinearity in different cell types plays distinct computational roles. Here we show that supralinearity in SST neurons, but not in other cell types, is important for explaining cotuned-ensemble suppression, showing that the role of supralinearity can be cell-type-dependent.

### Relation to previous theoretical works

Previous theoretical works on two-photon perturbation responses have typically relied on a linear approximation of the steady-state equations^8,21–23^. For modeling single-cell perturbations^21–23^, a linear approximation, which is equivalent to the quasi-linear approximation up to a scalar multiple, may be sufficient. Furthermore, a linear approximation can explain certain aspects of ensemble perturbation responses, such as compact cotuned ensemble perturbations recruiting more suppression than diffuse cotuned ensemble perturbations (as shown in Oldenburg et al. ^8^), a finding which we also reproduced. However, we found that a linear approximation cannot explain cotuned-ensemble suppression, where compact cotuned ensemble perturbations recruit more suppression than compact untuned ensemble perturbations. We showed that this effect is non-linear, such that a nonlinear theory that goes beyond a simple linear approximation is required for explaining these perturbation responses.

Unlike other previous works and this paper, Kong et al. ^23^ developed theory for understanding perturbation responses in ferret V1 rather than mouse V1. They found that single-cell perturbation responses in ferret V1 exhibit the same qualitative dependence on visual contrast as that predicted by earlier theoretical work in mouse V1^22^, suggesting that single-cell perturbation responses in ferret and mouse V1 may obey similar principles. However, unlike mouse V1, which has largely random (salt-and-pepper) arrangements of preferred orientation^71,72^, the organization of orientation tuning preferences in ferret V1 is quasi-periodic^73,74^. This may result in significant differences in tuned ensemble perturbation responses between mouse and ferret V1. For example, the salt-and-pepper organization of orientation tuning preferences in mouse V1 is crucial to our argument that cotuned-ensemble suppression is a nonlinear effect. It is possible that cotuned-ensemble suppression, if it exists in ferret V1, may be well accounted for by a linear model.

### Relation to holography in other brain areas

Optogenetic perturbations have emerged as a powerful tool for dissecting circuit function across multiple cortical areas, and our framework provides a principled approach to interpreting cell-type-specific responses to holographic perturbations in mouse V1. Beyond mouse V1, our framework can also be applied to study the spatial structure of perturbation responses in other cortical areas where temporal dynamics can be neglected. A natural next step is to incorporate the temporal dynamics of perturbation responses, which can reveal additional recurrent circuit mechanisms beyond those apparent at steady-state. For example, early responses may reflect monosynaptic contributions, while late responses may reflect polysynaptic contributions^75^. In motor cortex, the quick onset and offset of perturbation responses have been shown to be consistent with non-normal dynamics of E-I balanced networks with a low-dimensional task dynamics subspace^20^. Extending our framework to fit the temporal dynamics of perturbation responses, building on approaches previously developed in motor cortex^6^, promises to uncover further insights about the circuit mechanisms underlying cotuned perturbations.

### Limitations and future work

Our specific parametrization of the recurrent connectivity as a function of spatial distance, orientation tuning preference, and orientation selectivity is based on previous work^22^ and allows the steady-state perturbation response to be efficiently computed during model fitting. However, it imposes a strict constraint on the recurrent connectivity that is not fully consistent with experimental measurements. For example, our parametrization neglects preferential connections between neurons that are co-oriented and co-axially aligned^76^, or the fact that excitatory to excitatory connectivity is far more predictable from strong signal correlations than from common preferred orientation^77^. Furthermore, this limits the applicability of our approach to other cortical areas, for which this parametrization may not be suitable. Investigating perturbation responses in more flexible modeling frameworks such as low-rank RNNs^78–82^ may therefore be an interesting future direction.

Although we have included many biological constraints in our models to ensure consistency with various sources of experimental data, there are biological details not incorporated in our model that may play an important role in perturbation responses. For example, we did not model short-term plasticity at the synapses between different cell types, in particular the short-term facilitation of Pyr synapses onto SST neurons^83,84^. Future work incorporating these mechanisms could refine the picture of cell-type-specific contributions to perturbation responses.

## Conclusion

Understanding recurrent computations in the cortex is an outstanding challenge in neuroscience. We previously demonstrated an exactly solvable model with space- and feature-dependent connectivity, and used it to understand the circuitry implied by responses to single-cell perturbations^22^. Here we develop a systematic approach to understanding nonlinear aspects of steady state perturbation responses, and apply this to understand nonlinear aspects of responses to perturbations of small ensembles of neurons. By demonstrating a nonlinear inhibition pathway mediated by SST neurons that underlies these responses, we unveil a mechanism that may play an important role more generally in recurrent cortical computations. Our results highlight the power that analytic approaches can bring to wresting biological insights from responses to holographic optogenetic perturbations.

## RESOURCE AVAILABILITY

### Lead contact

Requests for further information and resources should be directed to and will be fulfilled by the lead contact, Ho Yin Chau.

### Materials availability

This study did not generate new materials.

### Data and code availability

- This paper analyzes existing, publicly available data, accessible at https://github.com/gregoryhandy/Logic_of_Recurrent_Circuits.
- All original code has been deposited at Zenodo and is publicly available at https://doi.org/10.5281/zenodo.22086790 as of the date of publication.
- Any additional information required to reanalyze the data reported in this paper is available from the lead contact upon request.

## ACKNOWLEDGMENTS

This work was supported by the Gatsby Charitable Foundation (GAT3708; and GAT3850 to A.P.), the Kavli Foundation, the NSF (DBI-1707398; and DGE-2036197 to H.Y.C.), the NIH (U01 NS108683, R01 EY029999, U19 NS107613; and T32 EY013933 to H.Y.C.), and the Simons

Foundation (1156607, to A.P.).

## AUTHOR CONTRIBUTIONS

Conceptualization, H.Y.C., K.D.M., and A.P.; methodology, H.Y.C., K.D.M., and A.P.; data curation, I.A.O. and H.Y.C.; formal analysis, H.Y.C.; investigation, H.Y.C.; writing–original draft, H.Y.C.; writing–review & editing, H.Y.C., K.D.M., and A.P.; funding acquisition, K.D.M. and A.P.; resources, K.D.M. and A.P.; supervision, K.D.M. and A.P.

## DECLARATION OF GENERATIVE AI AND AI-ASSISTED TECHNOLOGIES

During the preparation of this work, the authors used ChatGPT and Claude in order to review the manuscript for errors and suggest wording changes. After using this tool or service, the authors reviewed and edited the content as needed and take full responsibility for the content of the publication.

## SUPPLEMENTAL INFORMATION INDEX

Supplemental text, figures S1-S6, and their legends in a PDF

## METHODS

### Method details

#### Network models

The firing rate dynamics of all recurrent network models in this paper are given by

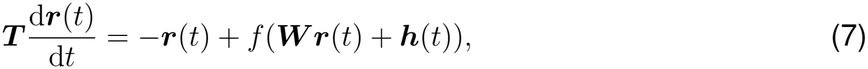

where ***r***(*t*) is the firing rate, ***W*** the connectivity matrix, ***h***(*t*) the external input, ***T*** a diagonal matrix of time constants, and *f* an arbitrary transfer function. ***h***(*t*) is equal to the baseline external input vector ***h*** for *t <* 0, and equal to ***h*** + *δ****h*** for *t* ≥ 0.

#### Network simulations

Network dynamics are simulated by solving the ordinary differential equation (ODE) Eq. 7 with an initial condition ***r***(0) that satisfies ***r***(0) = *f* (***W r***(0) + ***h***). The ODE is solved with Runge-Kutta methods of order 5(4), specifically with order-5 Dormand-Prince method for the random networks in Figure 1^85^ and a newer method by Tsitouras ^86^ for Figures 2-5. The random E-I networks in Figure 1 are simulated until *t* = 100*τ*_E_, where *τ*_E_ is the time constant of excitatory neurons, while the networks in Figures 2-5 are simulated until the network activity reaches steady-state. Steady-state is numerically determined by the condition |*∂*_*t*_*r*_*i*_| *≤ ϵ*_rel_ |*r*_*i*_|+ *ϵ*_abs_ for all neurons *i*, where *ϵ*_rel_ = 10^−4^, *ϵ*_abs_ = 5 × 10^−6^ for Figure 1, *ϵ*_rel_ = 5 × 10^−5^, *ϵ*_abs_ = 10^−6^ for Figures 2-3, and *ϵ*_rel_ = 5 × 10^−4^, *ϵ*_abs_ = 5 × 10^−6^ for Figures 4-5. For the networks in Figure 1, no output is returned if the neural activity at *t* = 100*τ*_E_ does not satisfy the steady-state condition, while for the networks in Figures 2-5, no output is returned if the neural activity fails to satisfy the steady-state condition within *t* = 200*τ*_E_.

#### Naive second-order approximation

The naive second order approximation for the steady-state perturbation response shown in Figure 1B is given by

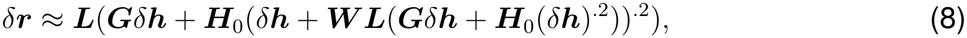

where 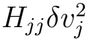. This is the approximation that results from Taylor expanding the nonlinearity *f* around ***v***^∗^ instead of ***v***^∗^ + *δ****h*** up to second order, such that it is exactly equal to the second-order approximation (Eq. 4) if the *n*-th derivative of *f* is 0 for all *n* > 2, *e*.*g. f* (*v*) = *v*^2^.

#### Nonlinear PV/SST/VIP pathways

Throughout Figures 4-5, when we refer to nonlinear PV/SST/VIP pathways (sometimes denoted (2,2,Y) where Y is one of PV, SST, or VIP), we specifically mean the sum

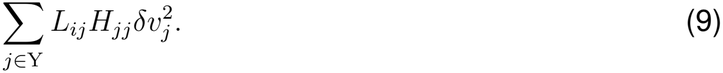

In Figure 4, *i* corresponds to a Pyr neuron, while in Figure 5, *i* corresponds to an inhibitory neuron with cell type X.

#### Random E-I networks

The random E-I networks in Figures 1 and S1 consist of 80% excitatory neurons and 20% inhibitory neurons (rounded to the nearest integer). Network weights are sampled from log-normal distributions, such that the linearized connectivity matrix ***GW*** has excitatory mean 1*/N*_*E*_, inhibitory mean *™*1.5*/N*_*I*_, and standard deviation 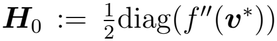 . All neurons have the same time constant. The transfer function of neurons is given by

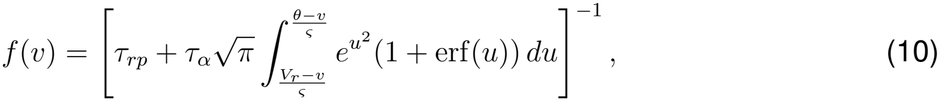

where *V*_*r*_ = 10 mV, *θ* = 20 mV, *ς* = 10 mV, *τ*_*rp*_ = 0.002 s, and *τ*_*α*_ = 0.02 s. Eq. 10 is the Ricciardi transfer function with parameters from Sanzeni et al. ^33^ . In our software implementation, the synaptic input *v* is represented in units of *ς* = 10 mV for convenience, such that the lower and upper bounds of the integral become 1 *™ v* and 2 *™ v* respectively. The integral in Eq. 10 is efficiently computed using a 5th order Gauss-Legendre quadrature rule, which achieves good numerical accuracy for all possible inputs *v*. The network’s firing rate is initialized at *r*_*i*_(0) = 1 Hz for all *i*, while the baseline external input vector ***h*** is given by *f*^−1^(***r***(0)) ***W r***(0), such that the initial firing rate ***r***(0) satisfies the unperturbed fixed-point equation ***r***(0) = *f* (***W r***(0) + ***h***). The perturbation vector *δ****h*** is given by *f*^−1^(***r***(0) + *δ****h***^*′*^) *f*^−1^(***r***(0)), where 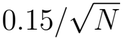 for *i* {1, · · ·, *N*_perturbed_ and zero otherwise, and where *δh*^*′*^ = 20 Hz for Figure 1B, while the values of *δh*^*′*^ used in Figure 1C are specified in the figure. *δh*^*′*^ is equal to the change in firing rate of the perturbed cells if no recurrent connections exist. We then compute the median relative error of the analytical approximations (Eq. 2, 3, 4, 8) compared to numerical simulations (details under “Network simulations” in Methods) across all unperturbed neurons. This procedure is repeated 300,000*/N* times to compute the median and CI of the median relative error across different random network instantiations.

#### Large-scale mouse V1 L2/3 network models

The network models in Figures 2-3 consist of 34,089 neurons located in a 1.15 mm × 1 mm 2D plane, while the models in Figures 4-5 consist of 40,759 neurons located in a 1.25 mm × 1.1 mm 2D plane. Each neuron in the network model is represented by the four-tuple (*α, µ*, ***x***, *θ*), where *α* is the cell type label, *µ* is a number between 0 and 1 representing orientation selectivity, ***x*** is a 2-dimensional vector representing spatial location, and *θ* is a circular variable between 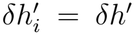 and 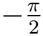representing orientation tuning preference.

The cell type label *α* is sampled from a categorical distribution with probabilities *p*_*α*_. For the models in Figures 2-3, *p*_E_ = 0.85, *p*_I_ = 0.15, while for models in Figures 4-5, *p*_Pyr_ = 0.85, *p*_PV_ = 0.043, *p*_SST_ = 0.032, *p*_VIP_ = 0.075. These values are chosen to roughly match the distribution of cell types in the biological model of mouse V1 L2/3 in Billeh et al. ^87^ . Orientation selectivity *µ* is sampled from a Beta distribution *B*(*a*_0_, *b*_0_) with *a*_0_ = 0.98, *b*_0_ = 1.28, where the parameters *a*_0_, *b*_0_ are obtained by fitting a Beta distribution to the empirical distribution of orientation selectivity in Oldenburg et al. ^8^ (Figure S2B). Spatial location ***x*** is sampled from a uniform distribution on the 2D simulation region. For each pair of neurons that are less than 3 µm apart, the spatial location of one of them is resampled until all pairwise distances between neurons are at least 3 µm. We do this because our parametrization of the connectivity strength between a pair of neurons, given by Eq. 11, tends to ∞ as the distance between them tends to 0, so this ensures that the connection strength cannot be arbitrarily large. Finally, the orientation tuning preference *θ* is sampled from a uniform distribution on 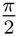.

Neurons are all-to-all connected, with the connection strength from neuron (*β, ν*, ***y***, *ϕ*) to neuron (*α, µ*, ***x***, *θ*) parametrized as

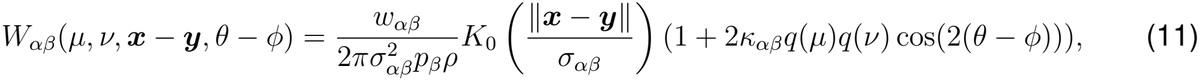

where *K*_0_ is the zeroth-order modified Bessel function of the second kind, *ρ* the neuronal density, *q* is a monotonically increasing function over the interval 0 to 1 with *q*(0) = 0 and *q*(1) = 1, and *w*_*αβ*_, *σ*_*αβ*_, *κ*_*αβ*_ are model parameters. In all models, the simulated cortical region and the number of neurons are chosen such that the neuronal density *ρ* is approximately the same as mouse V1 L2/3, with a value of ≈29,643 mm^−2 87^. Due to the lack of experimental constraints, *q* is chosen to be a function with theoretically convenient properties. Specifically, we define *q*(*µ*) = Φ(*µ*)^*p*^, where Φ is the cumulative distribution function of the distribution of orientation selectivity *µ* in the model, and *p* is chosen to be 0.25. This ensures *q* is monotonically increasing from *q*(0) = 0 to *q*(1) = 1, and that the quantity, E_*µ*_[*q*(*µ*)^*a*^], which is used in the analytical solution for the linearized perturbation response^22^, is simply (1 + *pa*)^−1^.

The time constants of all neurons are equal, except for the models in Figures 4-5, where the time constants of PV neurons are set to be half of that of Pyr neurons. The transfer function *f* is the Ricciardi nonlinearity (Eq. 10, with all parameters the same as in the random E-I networks except *τ*_*α*_ = 0.01 s for *α* = PV) for all neurons except for the Pyr neurons in Figures 4-5, which have instead a rectified linear transfer function, *f* (*v*) = *k*[*v*]_+_, where *k* = 0.1 Hz mV^−1^. Note that the exact value of *k* does not matter because any changes to *k* can be compensated by inversely proportional changes to the connection weights onto pyramidal neurons and the perturbation vector. A rectified linear transfer function is used for Pyr neurons in the models in Figures 4-5 because we found that the supralinearity of excitatory neurons has negligible effect on the perturbation responses (supralinearity of E cells only affects the (2,1) and (2,2,E) terms of the 2nd order approximation, and both terms are negligible as shown in Figure 3B-C), and that the perturbation response can be fitted to data much more efficiently when only the inhibitory cell types are supralinear (see explanation for Eq. 16, 17).

The network’s firing rate is initialized at ***r***(0) = *f* (***v***^∗^), while the baseline external input vector ***h*** is given by ***v***^∗^ − ***W*** *f* (***v***^∗^), where 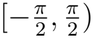 for all neurons *i* in the nonlinear models in Figures 2-3, and 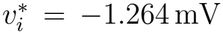 for Pyr neurons, −0.945 mV for PV, −0.921 mV for SST, and −0.334 mV for VIP in Figures 4-5. This ensures that the network is initialized at the unperturbed fixed point, *i*.*e*. ***r***(0) = *f* (***W r***(0) + ***h***). Furthermore, the values of ***v***^∗^ are chosen such that ***r***(0) is equal to the spontaneous firing rate values of 0.6 Hz for all neurons in Figures 2-3, and 0.6 Hz for Pyr, 1.35 Hz for PV, 0.682 Hz for SST, and 0.844 Hz for VIP in Figures 4-5. These cell-type specific spontaneous firing rate values are obtained from the publicly available code by Di Santo et al. ^68^, which in turn is obtained from the experimental data in Keller et al. ^45^ .

Finally, to roughly match the spatial distribution of experimentally recorded neurons (Figure S2A), only the responses of neurons within a 750 µm × 500 µm spatial window are analyzed. For the models in Figures 4-5, a bigger window of 950 µm × 800 µm is used for the inhibitory cell types to compensate for the small amount of neurons for each inhibitory cell type.

#### Ensemble perturbations

The ensemble perturbation vector *δ****h*** is defined as

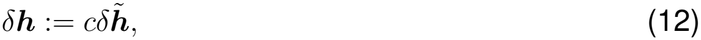

where *c* is a scalar parameter that is optimized, and 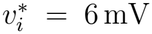 is a vector whose elements are either 0 or 1. The 10 perturbed cells (elements of 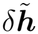 that are equal to 1) are selected with weighted random sampling (weighted sampling without replacement)^88^, where the weight of neuron *i* is given by

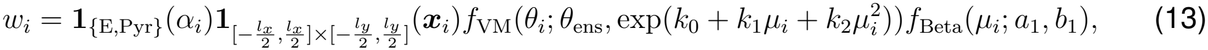

where *α*_*i*_, ***x***_*i*_, *θ*_*i*_, *µ*_*i*_ are the cell type, spatial location, preferred orientation, and orientation selectivity of neuron *i* respectively, **1**_*A*_ is the indicator function of a set *A* (*i*.*e*. **1**_*A*_(*x*) = 1 if *x* ∈ *A* and 0 otherwise), *f*_VM_(*θ*; *θ*_0_, *κ*) ∝ exp(*κ* cos(2(*θ*−*θ*_0_))) is the probability density function of the von Mises distribution, *f*_Beta_(*µ*; *a, b*) ∝ *µ*^*a*−1^(1 − *µ*)^*b*−1^ is the probability density function of the Beta distribution, *θ*_ens_ is a random sample from the uniform distribution 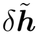 that is roughly equal to the ensemble tuning preference, and *l*_*x*_, *l*_*y*_, *k*_0_, *k*_1_, *k*_2_, *a*_1_, *b*_1_ are ensemble-specific parameters. For diffuse ensembles, *l*_*x*_ = 504.9 µm, *l*_*y*_ = 379.5 µm, while for compact ensembles, *l*_*x*_ = 292.4 µm, *l*_*y*_ = 258.3 µm. These parameter values are chosen such that the distribution of pairwise distances between perturbed cells approximates that of the diffuse and compact ensembles in Oldenburg et al. ^8^ (Figure S2E). For untuned ensembles, *k*_0_ = *™* 3.7, *k*_1_ = 10.0, *k*_2_ = *™* 9.8, *a*_1_ = 0.95, *b*_1_ = 1.22, while for cotuned ensembles, *k*_0_ = *™* 7.3, *k*_1_ = 19.0, *k*_2_ = *™* 12.0, *a*_1_ = 1.32, *b*_1_ = 0.66. These parameter values are obtained by fitting *f*_VM_(*θ*; *θ*_ens_, exp(*k*_0_ +*k*_1_*µ*+*k*_2_*µ*^2^))*f*_Beta_(*µ*; *a*_0_ +*a*_1_ *™* 1, *b*_0_ +*b*_1_ *™* 1) to the empirical distributions of orientation tuning preference and selectivity of the untuned and cotuned ensembles in Oldenburg et al. ^8^ respectively (Figure S2F). Finally, for untuned ensembles, we resample the 10 perturbed cells if the ensemble orientation selectivity, defined as 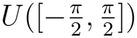 where *P* is the set of indices of perturbed cells, is greater than 0.07. This procedure is needed because our sampling method introduces a bias towards more cotuned ensembles, and this bias is particularly large for the sampling of untuned ensembles (Figure S2G).

#### Accounting for finite-size effect in linear models

We have shown theoretically that a linear model of mouse V1, whose connectivity is translationally-invariant in tuning preference and independent of presynaptic orientation selectivity when averaged over tuning preferences, cannot produce cotuned-ensemble suppression when the number of neurons is sufficiently large. However, a linear model with a sufficiently small number of neurons can produce cotuned-ensemble suppression through a finite-size effect (see supplemental text). While this effect is negligible in a network with realistic neuronal density, the same finite-size effect may still occur in the data due to the fact that only ∼1000 neurons are recorded and selected for ensemble perturbations^8^. To account for this potential effect, in the linear models in Figure 2, only 1000 neurons within the 750 µm × 500 µm spatial window were randomly selected to be “recorded,” and the perturbed ensembles were selected from these recorded neurons. This random selection was omitted in all other models since the finite-size effect was negligible in these linear models.

#### Fitting large-scale models

A naive approach for fitting steady-state perturbation responses to data is to simulate network dynamics numerically until steady-state, then backpropagate the error through time to update the model parameters. However, this approach can be prohibitively slow when many simulation steps are required. We address this issue by instead computing the steady-state perturbation response using an efficient semi-analytical approach. This approach combines the exact analytical solution for the perturbation response in a linear network we derived previously^22^ with an efficient fixed-point iteration scheme. Specifically, we initialize the two vectors

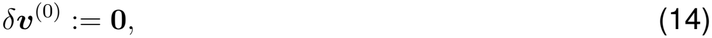

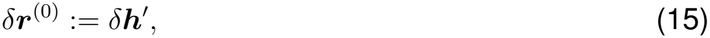

and then compute *δ****r***^(*n*)^ iteratively using the recurrence relations

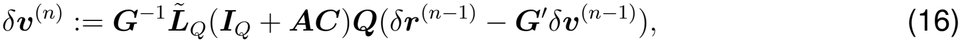

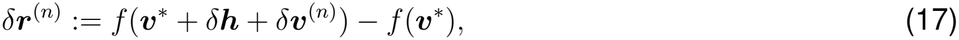

where *Q* is the number of cells that are either perturbed or have a nonlinear transfer function, ***I***_*Q*_ is the *Q* × *Q* identity matrix, and 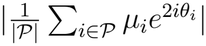 are defined in the supplemental text in which we derive Eq. 16, 17. Note that here *δ****v***^(*n*)^ is an iteration variable that converges to the exact change in recurrent input, not the first-order quantity *δ****v*** defined in Eq. 5. This fixed-point iteration approach has space complexity *O*(*NQ*), as opposed to *O*(*N*^2^) in a naive implementation. Since the number of perturbed cells is negligible, *Q* is approximately the number of cells with a nonlinear transfer function. As such, the perturbation response can be computed much more efficiently when excitatory cells, which are the majority of cells in the model, are assumed to be linear. In practice, we also assume that the perturbation response is weak enough that neurons with rectified linear transfer functions are effectively linear, allowing for efficient model fitting. Importantly, we find that 10 iterations of the recurrence relations (*n* = 10) are usually sufficient to produce accurate approximations of the perturbation response. As such, we always perform 10 iterations, unless the recurrence relations diverge, in which case we preemptively stop the process and return the last iteration.

The models are fitted to the perturbation response as a function of distance for all four types of ensemble perturbations (Figure S2C) as well as the perturbation response as a function of distance and tuning preference (Figure S2D) for both types of cotuned ensemble perturbations. Neurons in the model are binned in the same way as the data (15 µm bins in distance and 45^°^ bins in preferred orientation) when computing the loss. We assume that ΔF/F is a linear function of firing rate, such that the experimental measurement of the change in ΔF/F is equal to the model’s perturbation response *δ****r*** multiplied by a scalar parameter *γ*. A weighted root-mean-square loss is computed where each experimental data point is weighted inversely proportional to the square of its standard error. Data points in Figure S2D are further weighted by a factor of 0.25 since we want the fitted models to capture the cotuned-ensemble suppression effect (which is captured by the data points in Figure S2C) as well as possible. For the models in Figures 4-5, we include a regularization term in the loss function that penalizes large perturbation responses far away from the perturbation sites. Specifically, responses of Pyr cells beyond 250 µm from the nearest perturbed cell and the responses of PV, SST, and VIP cells beyond 325 µm are regularized. The full specification of the loss function used for training the models is given in the supplemental text (Eq. S61).

During optimization, the loss is computed for the mean perturbation response (approximated by the above fixed-point iteration method) across 5 random samples of each type of ensemble perturbation in a random instantiation of the network neurons (random sampling of neuronal cell type, spatial location, orientation tuning preference, and orientation selectivity). Since optimizing model parameters of a large nonlinear network is computationally expensive, smaller networks are used during optimization of the nonlinear models (24,838 neurons in a 0.98 mm × 0.855 mm 2D plane for the nonlinear models in Figures 2-3, and 34,089 neurons in a 1.15 mm × 1 mm 2D plane for the nonlinear models in Figures 4-5). After convergence, a validation loss is computed for the mean perturbation response (obtained by numerical simulations) across 5 random samples of each type of ensemble perturbation in 10 random instantiations of the network neurons (using the full-sized models). A fixed random seed is used for computing the validation loss so that the output is reproducible. If there is more than one random instantiation of the network neurons in which the numerical simulation fails to converge for any ensemble perturbation, the fitted model parameters are discarded. If the validation loss is greater than 0.75 (the loss is normalized such that if the perturbation response of every neuron is 0, then the loss is equal to 1), the fitted model parameters are also discarded. We repeat this procedure until we obtain 500 sets of fitted parameters for the models in Figures 2-3, and 5000 for the models in Figures 4-5.

The fitted parameters are *w*_*αβ*_, *σ*_*αβ*_, *κ*_*αβ*_ from Eq. 11, the perturbation strength *c* from Eq. 12, and the firing-rate-to-ΔF/F conversion factor *γ*. Note that in practice, we optimize the unitless quantity *g*_*α*_*w*_*αβ*_, where *g*_*α*_ is the gain of cell type *α* and is fixed by our choices of nonlinearity and spontaneous firing rates. For the linear model, *γ* can be absorbed into *c*, resulting in one fewer fitted parameter. Different constraints are imposed to ensure that the model parameters are biologically plausible and that the model is consistent with different experimental observations. Specifically we constrain: 1) the connections to be positive for E cells and negative for I cells (*w*_*α*E_ > 0, *w*_*α*I_ *<* 0), 2) the connection strength to not be too large (*g*_*α*_|*w*_*αβ*_| *<* 20), 3) the connection widths *σ*_*αβ*_ to be between 100 µm and 175 µm, 4) the connectivity tuning-dependence *κ*_*αβ*_ to be between *™*0.5 and 0.5 to ensure Dale’s law is satisfied, 5) the excitatory connections to be like-to-like^12,25,89,90^ (*κ*_EE_, *κ*_IE_ > 0 for Figures 2-3, *κ*_Pyr,Pyr_, *κ*_SST,Pyr_ > 0 for Figures 4-5), 6) the perturbation strength to be around 10 Hz^8^ (80 mV *< c <* 120 mV for models with linear or rectified linear E cells, and 9.422 mV *< c <* 11.510 mV for models with supralinear E cells, ensuring that 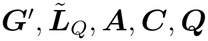 where *i* is a perturbed cell and *δ****h***^*′*^ is defined under Eq. 3), 7) the firing-rate-to-ΔF/F conversion factor *γ* to be upper bounded by the ratio of spontaneous ΔF/F (0.727 *±* 0.024) to spontaneous firing rate of Pyr cells (0.6 Hz) (0 *< γ <* 1.29 ≈ (0.727+2*0.024)*/*0.6 Hz^−1^); this constraint is combined with the constraint on *c* for the linear model, 8) the linearized network dynamics to be stable (see supplemental text for details), and 9) the network being inhibition-stabilized (*g*_E_*w*_EE_ > 1). For the models in Figures 4-5, we also constrain: 10) VIP-to-E, VIP-to-PV, VIP-to-VIP, SST-to-SST connections to be weak (*g*_*α*_ |*w*_*αβ*_| *<* 1), PV-to-SST connections to be not too strong (*g*_*α*_ *w*_*αβ*_ *<* 5), and reciprocal connections between SST and VIP neurons to be strong (2*g*_SST_| *w*_SST,VIP_ > *g*_Pyr_|*w*_Pyr,Pyr_|, 2*g*_VIP_ |*w*_VIP,SST_| > *g*_Pyr_|*w*_Pyr,Pyr_|)^16,84,87,91–93^, 11) the connection widths to depend only on presynaptic cell type identity in order to reduce the number of model parameters and improve the quality of fits (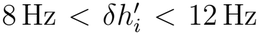 for all *α, α*^*′*^, *β*), 12) the reciprocal connections between Pyr and PV to be not strongly-tuned^94^ (|*κ*_Pyr,PV_|, |*κ*_PV,Pyr_| *<* 0.1), 13) PV neurons to be paradoxically suppressed by a weak activating input^17^ (*L*_PV,PV_ *<* 0), 14) SST neurons to *not* be paradoxically suppressed by a weak activating input, as they are unnecessary for stabilizing cortical activity at low stimulus contrast^41^ (*L*_SST,SST_ > 0), 15) responses of PV neurons to perturbation of Pyr neurons to be locally tuning-independent^12^ 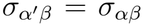, where 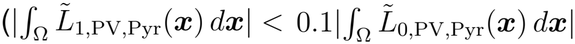 is given by Eq. S56, and Ω = {***x*** ∈ ℝ^2^ | ∥***x***∥_∞_ *<* 250 µm}), 16) responses of Pyr neurons to perturbation of SST neurons to be locally tuning-dependent^12^ 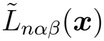, 17) Pyr neurons to be locally suppressed by perturbation of SST neurons^25^ (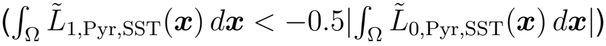, where 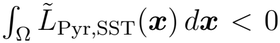 is given by Eq. S70), 18) Pyr neurons to be locally disinhibited by perturbation of VIP neurons ^25,26^ 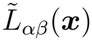, and 19) SST neurons to be strongly locally activated by perturbation of Pyr neurons^11^ (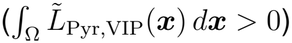 for all ∥***x***∥ _2_ ≤150 µm). The model parameters are initialized as follows: 1) *g*_*α*_*w*_*αβ*_ is sampled from a truncated normal distribution with standard deviation 0.5 and bounds according to the constraints above, 2) *σ*_*αβ*_, *κ*_*αβ*_ are sampled from a uniform distribution with bounds according to the constraints above, 3) *c* is initialized at 100 mV (*c* = 10.523 mV for models with supralinear E cells), and 4) *γ* is initialized at 1. Constrained optimization is performed using SLSQP (Sequential Least SQuares Programming)^95^ implemented in SciPy^96^, with gradients computed using PyTorch’s automatic differentiation engine^97^.

#### Difference in preferred orientation

In Figures 3F, 4E-G, S2D, and S3C, difference in preferred orientation from the perturbed ensembles (Δ preferred ori.) is defined as 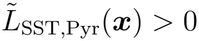, where *θ* and *θ*_ens_ are the preferred orientations of the measured cell and the perturbed ensemble respectively. Δ preferred ori. is then converted to degrees and binned into [0^°^, 22.5^°^), [22.5^°^, 67.5^°^), and [67.5^°^, 90^°^) bins, which are labeled as 0^°^, 45^°^, and 90^°^ respectively. In the data, the preferred orientation of the perturbed ensemble *θ*_ens_ is defined in the same way as the original paper^8^, *i*.*e. θ*_ens_ is the orientation of the drifting grating which elicited the maximum mean response. In our models, since we do not explicitly model visual responses, we instead define *θ*_ens_ as the weighted circular mean of the preferred orientations of cells in the perturbed ensemble, *i*.*e*. 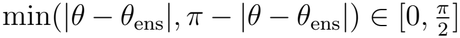 where *P* is the set of indices of perturbed cells and *θ*_*i*_, *µ*_*i*_ are the preferred orientation and orientation selectivity of cell *i* respectively.

### Data analysis

All data shown in this paper is from Oldenburg et al. ^8^ . Analysis of perturbation responses (Figures 2D-E, 4B, and S2C-D) follows the exact same procedure as Oldenburg et al. ^8^, with the only difference being that the perturbation responses are plotted against distance within the two-dimensional XY plane (lateral cortical distance) instead of the full three-dimensional XYZ space. This ensures consistency between the data analysis procedure and our model, which is two-dimensional in space. A brief summary of key analysis details is as follows (see Oldenburg et al. ^8^ for full details): ΔF/F is defined as (*F ™ F*_0_)*/F*_0_, where *F* is the neuropil-subtracted fluorescence, *F*_0_ is the rolling average of the 10^th^ percentile *F* over a ∼3-minute window. Perturbation response is measured as the difference between the mean ΔF/F 0.5 s to 1.5 s after the perturbation onset and the mean ΔF/F before the perturbation onset. Neuronal ensembles are classified as spatially compact if the mean pairwise distance between targeted neurons is *<* 200 µm, and classified as spatially diffuse otherwise. They are also classified as cotuned if the ensemble orientation selectivity index (OSI) is > 0.7 and the mean OSI of the perturbed cells is > 0.5, and untuned if the ensemble OSI is *<* 0.3 and the mean OSI is *<* 0.5.

In Figures S2B and S2F, parameters of the probability distributions are fitted with maximum likelihood estimation (MLE) using SciPy^96^.

### Quantification and statistical analysis

High-level descriptions of all statistical analyses performed are provided in the corresponding figure captions. Statistical tests were performed using SciPy^96^, while statistical visualizations were performed using seaborn^98^ and Matplotlib^99^.

All permutation tests were performed either exactly (described as an exact permutation test in the corresponding caption) or with 999,999 resamples. A permutation *t*-test refers to a permutation test using the *t*-statistic for the mean. Null distributions of the test statistic were generated by the following permutations: 1) Independent-samples permutation *t*-tests (Figure 2E) permute the ensemble-type labels (*e*.*g*. “Compact cotuned”) of all ensemble perturbations. 2) Paired permutation *t*-tests (Figures 2H, 2J, 3B, 3C, 4E-G, S2I, and S3A-B) permute the ensemble-type labels within each pair of ensemble perturbations, where two ensemble perturbations are paired if they are performed on the same random instantiation of a given model. 3) Permutation tests of Pearson correlation coefficient (Figures 5E, S5C-F, and S6) permute the y-values of all data points.

Exact Wilcoxon signed-rank tests (Figures 4C, 5C-D, and S5B) are exact paired permutation tests of the test statistic (signed-rank sums). The null distributions of the test statistic were generated by permuting the ensemble-type labels within each pair of mean responses, where two mean responses were paired if they were obtained from the same model.

