## Supplemental text and figures for "A nonlinear inhibition pathway underlying cortical responses to tuned holographic optogenetic perturbations"

Supplemental Information

Ho Yin Chau, Ian Antón Oldenburg, Kenneth D. Miller, and Agostina Palmigiano

2026

#### 1 Nonlinear theory of perturbation responses

We present here the derivation of Eqs. 2-4 in the main text. We consider a nonlinear rate-based RNN with $N$ neurons that follows typical rate dynamics (Eq. 7 in Methods). The connectivity matrix of the RNN is denoted $\boldsymbol{W} \in \mathbb{R}^{N \times N}$, and the nonlinear transfer function of the neurons is denoted $f : \mathbb{R} \to \mathbb{R}$. For any vector $\boldsymbol{v}$, we let $f(\boldsymbol{v})$ denote an elementwise application of $f$ to $\boldsymbol{v}$. Note that this derivation generalizes in a straightforward manner to the case where each neuron has a different transfer function, as in the 4-cell-type models in Figures 4-5.

Before the perturbation is applied, the network receives a baseline external input $\boldsymbol{h} \in \mathbb{R}^N$, and we assume that this input drives neural activity to a steady-state solution $\boldsymbol{r} \in \mathbb{R}^N$ that satisfies the fixed-point equation (Eq. 1 in the main text)

$$\boldsymbol{r} = f(\boldsymbol{v}^*) \tag{S1}$$

where $\boldsymbol{v}^* := \boldsymbol{W}\boldsymbol{r} + \boldsymbol{h}$. The network then receives a constant perturbation $\delta\boldsymbol{h} \in \mathbb{R}^N$, and we assume that this constant perturbation drives the network activity to a new steady-state solution. We assume the perturbation is sparse, *i.e.* $P \ll N$, where $P$ is the number of nonzero elements of $\delta\boldsymbol{h}$. The change in steady-state firing rate due to the perturbation is denoted $\delta\boldsymbol{r} \in \mathbb{R}^N$, and it satisfies the new fixed-point equation

$$\boldsymbol{r} + \delta\boldsymbol{r} = f(\boldsymbol{v}^* + \boldsymbol{W}\delta\boldsymbol{r} + \delta\boldsymbol{h}). \tag{S2}$$

To obtain the linear approximation (Eq. 2 in the main text), we Taylor expand $f$ in Eq. S2 around $\boldsymbol{v}^*$ up to first order as

$$\boldsymbol{r} + \delta\boldsymbol{r} \approx f(\boldsymbol{v}^*) + \mathrm{diag}(f'(\boldsymbol{v}^*))(\boldsymbol{W}\delta\boldsymbol{r} + \delta\boldsymbol{h}). \tag{S3}$$

Let $\boldsymbol{G}$ be the diagonal matrix of neuronal gain before the perturbation and $\boldsymbol{L}$ be the linear response matrix, *i.e.*

$$\boldsymbol{G} := \mathrm{diag}(f'(\boldsymbol{v}^*)), \tag{S4}$$

$$\boldsymbol{L} := (\boldsymbol{I} - \boldsymbol{G}\boldsymbol{W})^{-1}. \tag{S5}$$

Plugging these definitions into Eq. S3 and rearranging, we obtain the linear approximation

$$\delta\boldsymbol{r} \approx \boldsymbol{L}\boldsymbol{G}\delta\boldsymbol{h}. \tag{S6}$$

However, as explained in the main text, the nonzero elements of $\delta\boldsymbol{h}$ are large for typical holographic optogenetic perturbations. Since we Taylor expanded $f$ in powers of both $\boldsymbol{W}\delta\boldsymbol{r}$ and $\delta\boldsymbol{h}$ (Eq. S3), this results in large errors in the linear approximation. We therefore introduce the quasi-linear approximation, which improves on the linear approximation by Taylor expanding $f$ in Eq. S2 around $\boldsymbol{v}^* + \delta\boldsymbol{h}$ instead of $\boldsymbol{v}^*$, so that the Taylor expansion only involves powers of $\boldsymbol{W}\delta\boldsymbol{r}$. Performing the new Taylor expansion up to first order yields

$$\boldsymbol{r} + \delta\boldsymbol{r} \approx f(\boldsymbol{v}^* + \delta\boldsymbol{h}) + \mathrm{diag}(f'(\boldsymbol{v}^* + \delta\boldsymbol{h}))\boldsymbol{W}\delta\boldsymbol{r}. \tag{S7}$$

Next, we define

$$\delta\boldsymbol{h}' := f(\boldsymbol{v}^* + \delta\boldsymbol{h}) - \boldsymbol{r}, \tag{S8}$$

$$\delta\boldsymbol{G} := \mathrm{diag}(f'(\boldsymbol{v}^* + \delta\boldsymbol{h})) - \boldsymbol{G}, \tag{S9}$$

such that $\delta\boldsymbol{h}'$ and $\delta\boldsymbol{G}$ represent respectively the change in firing rate and change in neuronal gain due to the perturbation in the absence of recurrent connections. Plugging these definitions into Eq. S7 and rearranging, we obtain

$$\begin{aligned}\delta\boldsymbol{r} &\approx ((\boldsymbol{I} - \boldsymbol{G}\boldsymbol{W}) - \delta\boldsymbol{G}\boldsymbol{W})^{-1}\delta\boldsymbol{h}' \\ &= \boldsymbol{L}(\boldsymbol{I} - \delta\boldsymbol{G}\boldsymbol{W}\boldsymbol{L})^{-1}\delta\boldsymbol{h}'. \end{aligned} \tag{S10}$$

Next, notice that $\delta\boldsymbol{G}$ is a sparse diagonal matrix whose $i$-th diagonal element is zero for all $i \notin \mathcal{I}$, where $\mathcal{I}$ denotes the set of indices of perturbed cells. Also notice that $\boldsymbol{W}\boldsymbol{L} = -\boldsymbol{G}^{-1}(\boldsymbol{I} - \boldsymbol{G}\boldsymbol{W} - \boldsymbol{I})(\boldsymbol{I} - \boldsymbol{G}\boldsymbol{W})^{-1} = \boldsymbol{G}^{-1}(\boldsymbol{L} - \boldsymbol{I})$. We will assume that every element of $\tilde{\boldsymbol{L}} := \boldsymbol{L} - \boldsymbol{I}$ scales as $\mathcal{O}(N^{-1})$. This scaling assumption is satisfied when the connectivity matrix is obtained by discretizing a neural-field model and the elements of $\boldsymbol{W}$ scale as $\mathcal{O}(N^{-1})$[1], a common approach to modeling cortical connectivity[1,2]. With the scaling assumption of $\tilde{\boldsymbol{L}}$ and the sparsity of $\delta\boldsymbol{G}$, we see that $\delta\boldsymbol{G}\boldsymbol{W}\boldsymbol{L}$ is a matrix whose $i$-th row is zero for all $i \notin \mathcal{I}$, and every element of its nonzero rows scales as $\mathcal{O}(N^{-1})$. Thus

$$\|\delta\boldsymbol{G}\boldsymbol{W}\boldsymbol{L}\|_2 \leq \|\delta\boldsymbol{G}\boldsymbol{W}\boldsymbol{L}\|_F = \mathcal{O}(N^{-\frac{1}{2}}). \tag{S11}$$

Therefore, for sufficiently large $N$, we must have $\|\delta\boldsymbol{G}\boldsymbol{W}\boldsymbol{L}\|_2 < 1$, so we can expand the inverse $(\boldsymbol{I}-\delta\boldsymbol{G}\boldsymbol{W}\boldsymbol{L})^{-1}$ as a Neumann series and obtain

$$\delta\boldsymbol{r} \approx \boldsymbol{L}\sum_{n=0}^{\infty}(\delta\boldsymbol{G}\boldsymbol{W}\boldsymbol{L})^n\delta\boldsymbol{h}'. \tag{S12}$$

But now notice that for all $n \geq 1$, $(\delta\boldsymbol{G}\boldsymbol{W}\boldsymbol{L})^n$ is also a matrix whose $i$-th row is zero for all $i \notin \mathcal{I}$, and every element of its nonzero rows scales as $\mathcal{O}(N^{-n})$. Thus $(\delta\boldsymbol{G}\boldsymbol{W}\boldsymbol{L})^n\delta\boldsymbol{h}'$ is a sparse vector whose $i$-th element is zero for all $i \notin \mathcal{I}$ and whose nonzero elements scale as $\mathcal{O}(N^{-n})$, and $\tilde{\boldsymbol{L}}(\delta\boldsymbol{G}\boldsymbol{W}\boldsymbol{L})^n\delta\boldsymbol{h}'$ is a vector whose elements all scale as $\mathcal{O}(N^{-(n+1)})$. Letting $\boldsymbol{e}_i$ denote the $i$-th unit vector, we therefore have

$$\boldsymbol{e}_i^T\boldsymbol{L}(\delta\boldsymbol{G}\boldsymbol{W}\boldsymbol{L})^n\delta\boldsymbol{h}' = \begin{cases}\mathcal{O}(N^{-(n+1)}), & i \notin \mathcal{I} \\ \mathcal{O}(N^{-n}), & \text{otherwise}\end{cases} \tag{S13}$$

for all $n \geq 0$. Thus, if we consider only the lowest order term ($\mathcal{O}(N^{-1})$ for the unperturbed cells), we obtain the quasi-linear approximation

$$\delta \boldsymbol{r} \approx \boldsymbol{L}\delta \boldsymbol{h}'. \tag{S14}$$

To obtain the second order approximation, we Taylor expand $f$ in Eq. S2 around $\boldsymbol{v}^* + \delta \boldsymbol{h}$ up to second order. This yields

$$\boldsymbol{r} + \delta \boldsymbol{r} \approx f(\boldsymbol{v}^* + \delta \boldsymbol{h}) + \mathrm{diag}(f'(\boldsymbol{v}^* + \delta \boldsymbol{h}))\boldsymbol{W}\delta \boldsymbol{r} + \frac{1}{2}\mathrm{diag}(f''(\boldsymbol{v}^* + \delta \boldsymbol{h}))(\boldsymbol{W}\delta \boldsymbol{r})^{\cdot 2}. \tag{S15}$$

Next, we define

$$\boldsymbol{H} := \frac{1}{2}\mathrm{diag}(f''(\boldsymbol{v}^* + \delta \boldsymbol{h})). \tag{S16}$$

Then Eq. S15 can be simplified as

$$\begin{aligned}\delta \boldsymbol{r} &\approx ((\boldsymbol{I} - \boldsymbol{G}\boldsymbol{W}) - \delta \boldsymbol{G}\boldsymbol{W})^{-1}(\delta \boldsymbol{h}' + \boldsymbol{H}(\boldsymbol{W}\delta \boldsymbol{r})^{\cdot 2}) \\ &= \boldsymbol{L}(\boldsymbol{I} - \delta \boldsymbol{G}\boldsymbol{W}\boldsymbol{L})^{-1}(\delta \boldsymbol{h}' + \boldsymbol{H}(\boldsymbol{W}\delta \boldsymbol{r})^{\cdot 2})\end{aligned} \tag{S17}$$

Again, assuming $\tilde{\boldsymbol{L}}$ scales as $\mathcal{O}(N^{-1})$ and that $N$ is sufficiently large, we can expand the inverse $(\boldsymbol{I} - \delta \boldsymbol{G}\boldsymbol{W}\boldsymbol{L})^{-1}$ as a Neumann series, yielding

$$\delta \boldsymbol{r} \approx \boldsymbol{L}\sum_{n=0}^{\infty}(\delta \boldsymbol{G}\boldsymbol{W}\boldsymbol{L})^n(\delta \boldsymbol{h}' + \boldsymbol{H}(\boldsymbol{W}\delta \boldsymbol{r})^{\cdot 2}). \tag{S18}$$

To arrive at the second order approximation, we plug in the quasi-linear approximation of $\delta \boldsymbol{r}$ into the right hand side of Eq. S18, yielding

$$\delta \boldsymbol{r} \approx \boldsymbol{L}\sum_{n=0}^{\infty}(\delta \boldsymbol{G}\boldsymbol{W}\boldsymbol{L})^n(\delta \boldsymbol{h}' + \boldsymbol{H}(\boldsymbol{W}\boldsymbol{L}\delta \boldsymbol{h}')^{\cdot 2}). \tag{S19}$$

Since $\boldsymbol{W}\boldsymbol{L} = \boldsymbol{G}^{-1}\tilde{\boldsymbol{L}}$, every element of $\boldsymbol{H}(\boldsymbol{W}\boldsymbol{L}\delta \boldsymbol{h}')^{\cdot 2}$ scales as $\mathcal{O}(N^{-2})$. Thus for all $n \geq 1$, $(\delta \boldsymbol{G}\boldsymbol{W}\boldsymbol{L})^n\boldsymbol{H}(\boldsymbol{W}\boldsymbol{L}\delta \boldsymbol{h}')^{\cdot 2}$ is a sparse vector whose $i$-th element is zero for all $i \notin \mathcal{I}$ and whose nonzero elements scale as $\mathcal{O}(N^{-(n+1)})$, and $\tilde{\boldsymbol{L}}(\delta \boldsymbol{G}\boldsymbol{W}\boldsymbol{L})^n\boldsymbol{H}(\boldsymbol{W}\boldsymbol{L}\delta \boldsymbol{h}')^{\cdot 2}$ is a vector whose elements all scale as $\mathcal{O}(N^{-(n+2)})$. Thus

$$\boldsymbol{e}_i^T\boldsymbol{L}(\delta \boldsymbol{G}\boldsymbol{W}\boldsymbol{L})^n\boldsymbol{H}(\boldsymbol{W}\boldsymbol{L}\delta \boldsymbol{h}')^{\cdot 2} = \begin{cases}\mathcal{O}(N^{-(n+2)}), & i \notin \mathcal{I} \\ \mathcal{O}(N^{-(n+1)}), & \text{otherwise}\end{cases} \tag{S20}$$

for all $n \geq 1$. Collecting all terms up to $\mathcal{O}(N^{-2})$ for the unperturbed cells (the order of each term can be determined from Eq. S13 and Eq. S20), we have the second-order approximation

$$\delta \boldsymbol{r} \approx \boldsymbol{L}\delta \boldsymbol{h}' + \boldsymbol{L}\delta \boldsymbol{G}\boldsymbol{W}\boldsymbol{L}\delta \boldsymbol{h}' + \boldsymbol{L}\boldsymbol{H}(\boldsymbol{W}\boldsymbol{L}\delta \boldsymbol{h}')^{\cdot 2}. \tag{S21}$$

Defining $\delta \boldsymbol{v} := \boldsymbol{W}\boldsymbol{L}\delta \boldsymbol{h}'$ and plugging it into Eq. S21 then yields Eq. 4 in the main text.

#### 2 Cotuned-ensemble suppression is a nonlinear effect

Here we provide a more formal justification that a linear model with a sufficiently large number of neurons, assuming that its connectivity 1) depends on orientation tuning preference only through the difference between pre- and post-synaptic tuning preferences (translation symmetry), and 2) is independent of presynaptic orientation selectivity when averaged over tuning preferences (selectivity independence), cannot explain cotuned-ensemble suppression in mouse V1. Note that the connectivity of our models (Eq. 11 in Methods) satisfies both assumptions. To show this, we start with the equation defining the steady-state perturbation response in a linear model, given by

$$\delta r_i = \sum_{j=1}^{N} W_{ij}\delta r_j + \delta h_i. \tag{S22}$$

for all $i$. We will assume that each neuron $i$ is uniquely characterized by its orientation tuning preference $\theta_i$, orientation selectivity $\mu_i$, and some vector $\boldsymbol{x}_i$ (for example $\boldsymbol{x}_i$ could represent the spatial location of neuron $i$). Then Eq. S22 can be written as

$$\delta r(\theta_i, \mu_i, \boldsymbol{x}_i) = \sum_{j=1}^{N} W(\theta_i - \theta_j, \mu_i, \mu_j, \boldsymbol{x}_i, \boldsymbol{x}_j)\delta r(\theta_j, \mu_j, \boldsymbol{x}_j) + \delta h(\theta_i, \mu_i, \boldsymbol{x}_i) \tag{S23}$$

for some appropriately-defined functions $\delta r, W, \delta h$. Note that $W$ only depends on orientation tuning preference through the difference $\theta_i - \theta_j$ by assumption 1, and that $W$ is $2\pi$-periodic in $\theta$ because orientation tuning preference is a circular variable, such that

$$\int_{-\pi}^{\pi} W(\theta - \phi, \mu, \nu, \boldsymbol{x}, \boldsymbol{y})\, d\theta = \int_{-\pi}^{\pi} W(\theta, \mu, \nu, \boldsymbol{x}, \boldsymbol{y})\, d\theta =: \langle W\rangle_\theta(\mu, \nu, \boldsymbol{x}, \boldsymbol{y}) \tag{S24}$$

for all $\phi, \mu, \nu, \boldsymbol{x}, \boldsymbol{y}$ (in this section we'll let orientation tuning preference be a variable in $[-\pi, \pi)$ instead of $[-\frac{\pi}{2}, \frac{\pi}{2})$ for mathematical convenience). Furthermore, by assumption 2, we have

$$\langle W\rangle_\theta(\mu, \nu, \boldsymbol{x}, \boldsymbol{y}) = \langle W\rangle_\theta(\mu, \boldsymbol{x}, \boldsymbol{y}). \tag{S25}$$

When the number of neurons is sufficiently large, we can take the continuum limit of Eq. S23 and obtain the field equation

$$\delta r(\theta, \mu, \boldsymbol{x}) = \int_{\Omega}\int_0^1\int_{-\pi}^{\pi} W(\theta - \phi, \mu, \nu, \boldsymbol{x}, \boldsymbol{y})\delta r(\phi, \nu, \boldsymbol{y})p(\nu)\, d\phi\, d\nu\, d\eta(\boldsymbol{y}) + \delta h(\theta, \mu, \boldsymbol{x}) \tag{S26}$$

for some integration region $\Omega$, measure $\eta$, and probability density function of orientation selectivity $p(\nu)$, with the additional assumptions that the distribution of orientation tuning preferences is uniform and that the distributions of $\theta, \mu, \boldsymbol{x}$ are independent. These assumptions are mostly consistent with mouse V1, which has a largely salt-and-pepper organization of orientation tuning preferences[3,4], although they ignore the fine spatial scale clustering of orientation tuning preferences as recently found by Yu et al.[5].

We now consider the response to an ensemble perturbation by defining

$$\delta h(\theta, \mu, \boldsymbol{x}) := \sum_{i=1}^{P} \frac{c_i}{p(\mu_i)}\delta(\theta - \theta_i)\delta(\mu - \mu_i)\delta(\boldsymbol{x} - \boldsymbol{x}_i) \tag{S27}$$

for some choice of $c_i, \theta_i, \mu_i, \boldsymbol{x}_i$ for $i \in \{1, \cdots, P\}$, where $P$ is the number of perturbed cells, and the prefactors $p^{-1}(\mu_i)$ ensure that the total synaptic input to the network due to the perturbation,

$$\int_\Omega \int_0^1 \int_{-\pi}^{\pi} \delta h(\theta, \mu, \boldsymbol{x}) p(\mu)\, d\theta\, d\mu\, d\eta(\boldsymbol{x}) \tag{S28}$$

is independent of the orientation selectivities $\mu_i$ of the perturbed cells. Our claim is that the response to an ensemble perturbation, when averaged over orientation tuning preference and selectivity, is independent of the choices of $\theta_i$ and $\mu_i$. More formally, we show that the integral,

$$\langle \delta r \rangle_{\theta,\mu}(\boldsymbol{x}) := \int_0^1 \int_{-\pi}^{\pi} \delta r(\theta, \mu, \boldsymbol{x}) p(\mu)\, d\theta\, d\mu, \tag{S29}$$

which is proportional to the mean response at $\boldsymbol{x}$, is defined by a self-consistency equation that has no dependence on $\theta_i$ or $\mu_i$. Indeed, if we apply the integral operator $\int_{-\pi}^{\pi} d\theta \int_0^1 d\mu\, p(\mu)$ to both sides of Eq. S26, we obtain

$$\langle \delta r \rangle_{\theta,\mu}(\boldsymbol{x}) = \int_\Omega \int_0^1 \int_{-\pi}^{\pi} \left( \int_0^1 \int_{-\pi}^{\pi} W(\theta - \phi, \mu, \nu, \boldsymbol{x}, \boldsymbol{y}) p(\mu)\, d\theta\, d\mu \right) \delta r(\phi, \nu, \boldsymbol{y}) p(\nu)\, d\phi\, d\nu\, d\eta(\boldsymbol{y}) + \sum_{i=1}^{P} c_i \delta(\boldsymbol{x} - \boldsymbol{x}_i). \tag{S30}$$

By Eq. S24 and Eq. S25, we have

$$\int_0^1 \int_{-\pi}^{\pi} W(\theta - \phi, \mu, \nu, \boldsymbol{x}, \boldsymbol{y}) p(\mu)\, d\theta\, d\mu = \int_0^1 \langle W \rangle_\theta(\mu, \boldsymbol{x}, \boldsymbol{y}) p(\mu) d\mu =: \langle W \rangle_{\theta,\mu}(\boldsymbol{x}, \boldsymbol{y}), \tag{S31}$$

so we have

$$\begin{aligned} \langle \delta r \rangle_{\theta,\mu}(\boldsymbol{x}) &= \int_\Omega \langle W \rangle_{\theta,\mu}(\boldsymbol{x}, \boldsymbol{y}) \left( \int_0^1 \int_{-\pi}^{\pi} \delta r(\phi, \nu, \boldsymbol{y}) p(\nu)\, d\phi\, d\nu \right) d\eta(\boldsymbol{y}) + \sum_{i=1}^{P} c_i \delta(\boldsymbol{x} - \boldsymbol{x}_i) \\ &= \int_\Omega \langle W \rangle_{\theta,\mu}(\boldsymbol{x}, \boldsymbol{y}) \langle \delta r \rangle_{\theta,\mu}(\boldsymbol{y})\, d\eta(\boldsymbol{y}) + \sum_{i=1}^{P} c_i \delta(\boldsymbol{x} - \boldsymbol{x}_i), \end{aligned}$$

which is, as claimed, a self-consistency equation for $\langle \delta r \rangle_{\theta,\mu}$ that doesn't depend on $\theta_i$ or $\mu_i$. Thus cotuned-ensemble suppression, under the assumptions prescribed, must be a nonlinear effect.

We note that as a corollary, the integral

$$\langle \delta r \rangle_\theta(\mu, \boldsymbol{x}) := \int_{-\pi}^{\pi} \delta r(\theta, \mu, \boldsymbol{x})\, d\theta, \tag{S32}$$

which is proportional to the mean response at $(\mu, \boldsymbol{x})$, is also independent of $\theta_i$ and $\mu_i$ when $(\mu, \boldsymbol{x}) \neq (\mu_i, \boldsymbol{x}_i)$ for all $i \in \{1, \cdots, P\}$. Indeed, if we apply the tuning preference integral $\int_{-\pi}^{\pi} d\theta$ to both sides of Eq. S26, we obtain

$$\langle \delta r \rangle_\theta(\mu, \boldsymbol{x}) = \int_\Omega \langle W \rangle_\theta(\mu, \boldsymbol{x}, \boldsymbol{y}) \langle \delta r \rangle_{\theta,\mu}(\boldsymbol{y})\, d\eta(\boldsymbol{y}) + \sum_{i=1}^{P} \frac{c_i}{p(\mu_i)} \delta(\mu - \mu_i) \delta(\boldsymbol{x} - \boldsymbol{x}_i). \tag{S33}$$

When $(\mu, \boldsymbol{x}) \neq (\mu_i, \boldsymbol{x}_i)$ for all $i \in \{1, \cdots, P\}$, the Dirac delta function term vanishes, and thus $\langle \delta r \rangle_\theta(\mu, \boldsymbol{x})$ is also independent of $\theta_i$ and $\mu_i$.

### 3 Finite-size effect in linear models

To show that cotuned-ensemble suppression is a nonlinear effect, we assumed that the number of neurons is sufficiently large, such that we can take the continuum limit of the perturbation response. This assumption may be violated in the experimental data, since only ∼1000 neurons are recorded and selected for ensemble perturbations in each session[6]. As a result, a linear model may potentially be able to explain cotuned-ensemble suppression through a finite-size effect. To illustrate this effect, consider a linear network with only 20 E cells, half of which are tuned to 0° stimuli, while the other half are tuned to 90° stimuli. Suppose connectivity in this linear model is tuned in such a way that perturbing an E cell excites similarly-tuned E cells but suppresses oppositely-tuned E cells by the same amount. A cotuned ensemble perturbation of all 10 0° E cells would strongly suppress the 10 90° E cells. But since these 90° E cells are the only unperturbed cells, the average perturbation response (which excludes all perturbed cells) would be strongly suppressed. On the other hand, an untuned ensemble perturbation of 5 0° and 5 90° E cells would result in net zero input to the remaining E cells, causing the average perturbation response to be roughly 0. Thus cotuned-ensemble suppression will be observed in this linear model. As explained in STAR Methods, we account for this potential finite-size effect when fitting our linear models by selecting only 1000 neurons in our model to be "recorded" and chosen for ensemble perturbations. The fact that our fitted linear models fail to exhibit cotuned-ensemble suppression suggests that this finite-size effect is negligible in the experimental data.

### 4 Sign of $L_{ij}$

Here we expand on the brief remark we made in the main text that in our E-I models, $L_{ij}$, the elements of the linear response matrix $\boldsymbol{L}$, are negative when $j$ is an I cell and $L_{ij}$ is averaged across all E cells $i$. In other words, there are no I cells which cause disinhibition of E cells on average in our E-I models. This assumes the number of neurons is large enough that $L_{ij}$ can be well described by the exact analytical solution for the linear perturbation response we previously developed[1]. Indeed, by Eq. S55 in the supplemental text of Chau et al.[1], we have

$$\langle L_{ij} \rangle_{i \in \alpha} = N_\beta^{-1} [(\boldsymbol{I} - \boldsymbol{J})^{-1}]_{\alpha\beta}, \tag{S34}$$

where $\beta$ is the cell type of $j$, $N_\beta$ is the number of neurons with cell type $\beta$, and $\boldsymbol{J}$ is the $2 \times 2$ matrix

$$\boldsymbol{J} := \begin{bmatrix} g_{\mathrm{E}} w_{\mathrm{EE}} & g_{\mathrm{E}} w_{\mathrm{EI}} \\ g_{\mathrm{I}} w_{\mathrm{IE}} & g_{\mathrm{I}} w_{\mathrm{II}} \end{bmatrix}, \tag{S35}$$

and $g_\alpha$ is the neuronal gain of cell type $\alpha$. But now

$$[(\boldsymbol{I} - \boldsymbol{J})^{-1}]_{\mathrm{EI}} = \frac{g_{\mathrm{E}} w_{\mathrm{EI}}}{\det(\boldsymbol{I} - \boldsymbol{J})}, \tag{S36}$$

and $\det(\boldsymbol{I} - \boldsymbol{J})$ is positive for all linearly stable networks (Eq. S79 in the supplemental text of Chau et al.[1]), so since $w_{\mathrm{EI}} < 0$, $[(\boldsymbol{I} - \boldsymbol{J})^{-1}]_{\mathrm{EI}}$ is also negative, proving our claim.

### 5 Fitting large-scale models

#### 5.1 Derivation of recurrence relations for steady-state computation

Here we derive the recurrence relations for $\delta\boldsymbol{v}^{(n)}, \delta\boldsymbol{r}^{(n)}$ with which we approximately compute the steady-state perturbation response efficiently. We start by defining the function

$$\varrho(\boldsymbol{x}) := f(\boldsymbol{v}^* + \delta\boldsymbol{h} + \boldsymbol{x}) - \operatorname{diag}(f'(\boldsymbol{v}^* + \delta\boldsymbol{h}))\boldsymbol{x} - f(\boldsymbol{v}^* + \delta\boldsymbol{h}) \tag{S37}$$

such that by definition, the fixed-point equation Eq. S2 is exactly equal to

$$\boldsymbol{r} + \delta\boldsymbol{r} = f(\boldsymbol{v}^* + \delta\boldsymbol{h}) + \operatorname{diag}(f'(\boldsymbol{v}^* + \delta\boldsymbol{h}))\boldsymbol{W}\delta\boldsymbol{r} + \varrho(\boldsymbol{W}\delta\boldsymbol{r}). \tag{S38}$$

Note that this is just our second-order Taylor expansion (Eq. S15), but with the second-order term $\boldsymbol{H}(\boldsymbol{W}\delta\boldsymbol{r})^{\cdot 2}$ replaced with the exact residual $\varrho(\boldsymbol{W}\delta\boldsymbol{r})$. Rearranging in the same way as we did with Eq. S17, we obtain

$$\delta\boldsymbol{r} = \boldsymbol{L}(\boldsymbol{I} - \delta\boldsymbol{G}\boldsymbol{W}\boldsymbol{L})^{-1}(\delta\boldsymbol{h}' + \varrho(\boldsymbol{W}\delta\boldsymbol{r})). \tag{S39}$$

In principle we could simply perform fixed point iteration on this equation to compute $\delta\boldsymbol{r}$, but this approach is inefficient in several ways. First, this equation involves two dense $N \times N$ matrices $\boldsymbol{W}, \boldsymbol{L}$ ($\boldsymbol{G}$ is a diagonal matrix so it can be stored efficiently as a vector), which uses a lot of memory. Optimizing memory usage is paramount because $N$ is large in our models, backpropagation uses a lot of additional memory, and we perform all our computations on a single GPU, which has limited VRAM. Our first observation is that we only need to store a single dense $N \times N$ matrix, $\tilde{\boldsymbol{L}}$, if we left-multiply both sides by $\boldsymbol{W}$, define $\delta\tilde{\boldsymbol{v}} := \boldsymbol{W}\delta\boldsymbol{r}$, and note that $\boldsymbol{W}\boldsymbol{L} = \boldsymbol{G}^{-1}\tilde{\boldsymbol{L}}$, such that we obtain

$$\delta\tilde{\boldsymbol{v}} = \boldsymbol{G}^{-1}\tilde{\boldsymbol{L}}(\boldsymbol{I} - \delta\boldsymbol{G}\boldsymbol{G}^{-1}\tilde{\boldsymbol{L}})^{-1}(\delta\boldsymbol{h}' + \varrho(\delta\tilde{\boldsymbol{v}})). \tag{S40}$$

Second, this equation involves computing the large $N \times N$ matrix inverse $(\boldsymbol{I} - \delta\boldsymbol{G}\boldsymbol{G}^{-1}\tilde{\boldsymbol{L}})^{-1}$, which is inefficient. We can optimize this inverse by recalling that $\delta\boldsymbol{G}$ is a sparse diagonal matrix with only $P$ nonzero diagonal elements, where $P$ is the number of perturbed cells. As such, if we define the projection matrix

$$\boldsymbol{P} = \sum_{i=1}^{P} \boldsymbol{e}_{P,i}\boldsymbol{e}_{N,p_i}^T \in \mathbb{R}^{P \times N}, \tag{S41}$$

where $\{p_i\}_{i=1}^P$ are the indices of perturbed cells and $\boldsymbol{e}_{P,i}$ denotes the $i$-th unit vector in $\mathbb{R}^P$, then we have $\delta\boldsymbol{G} = \boldsymbol{P}^T\boldsymbol{P}\delta\boldsymbol{G}$. We can then apply the Woodbury matrix identity[7] to obtain an equivalent expression that only performs a small $P \times P$ matrix inverse,

$$(\boldsymbol{I} - \delta\boldsymbol{G}\boldsymbol{G}^{-1}\tilde{\boldsymbol{L}})^{-1} = (\boldsymbol{I} - \boldsymbol{P}^T\boldsymbol{P}\delta\boldsymbol{G}\boldsymbol{G}^{-1}\tilde{\boldsymbol{L}})^{-1} = \boldsymbol{I} + \boldsymbol{P}^T(\boldsymbol{I}_P - \boldsymbol{P}\delta\boldsymbol{G}\boldsymbol{G}^{-1}\tilde{\boldsymbol{L}}\boldsymbol{P}^T)^{-1}\boldsymbol{P}\delta\boldsymbol{G}\boldsymbol{G}^{-1}\tilde{\boldsymbol{L}}. \tag{S42}$$

Third, we notice that the vector $\delta\boldsymbol{h}' + \varrho(\delta\tilde{\boldsymbol{v}})$ only has $Q$ nonzero elements, where $Q$ is number of cells which are either perturbed or have a nonlinear transfer function. As such, if we define the projection matrix

$$\boldsymbol{Q} := \sum_{i=1}^{Q} \boldsymbol{e}_{Q,i}\boldsymbol{e}_{N,q_i}^T \in \mathbb{R}^{Q \times N}, \tag{S43}$$

where $\{q_i\}_{i=1}^Q$ are the indices of cells which are either perturbed or nonlinear, then $\delta \boldsymbol{h}' + \varrho(\delta\tilde{\boldsymbol{v}}) = \boldsymbol{Q}^T\boldsymbol{Q}(\delta\boldsymbol{h}' + \varrho(\delta\tilde{\boldsymbol{v}}))$. By similar reasoning, we also have $\boldsymbol{P}^T = \boldsymbol{Q}^T\boldsymbol{Q}\boldsymbol{P}^T$. Thus

$$\begin{aligned}\delta\tilde{\boldsymbol{v}} &= \boldsymbol{G}^{-1}\tilde{\boldsymbol{L}}(\boldsymbol{I} + \boldsymbol{Q}^T\boldsymbol{Q}\boldsymbol{P}^T(\boldsymbol{I}_P - \boldsymbol{P}\delta\boldsymbol{G}\boldsymbol{G}^{-1}\tilde{\boldsymbol{L}}\boldsymbol{Q}^T\boldsymbol{Q}\boldsymbol{P}^T)^{-1}\boldsymbol{P}\delta\boldsymbol{G}\boldsymbol{G}^{-1}\tilde{\boldsymbol{L}})\boldsymbol{Q}^T\boldsymbol{Q}(\delta\boldsymbol{h}' + \varrho(\delta\tilde{\boldsymbol{v}})) \\ &= \boldsymbol{G}^{-1}\tilde{\boldsymbol{L}}(\boldsymbol{Q}^T + \boldsymbol{Q}^T\boldsymbol{Q}\boldsymbol{P}^T(\boldsymbol{I}_P - \boldsymbol{P}\delta\boldsymbol{G}\boldsymbol{G}^{-1}\tilde{\boldsymbol{L}}\boldsymbol{Q}^T\boldsymbol{Q}\boldsymbol{P}^T)^{-1}\boldsymbol{P}\delta\boldsymbol{G}\boldsymbol{G}^{-1}\tilde{\boldsymbol{L}}\boldsymbol{Q}^T)\boldsymbol{Q}(\delta\boldsymbol{h}' + \varrho(\delta\tilde{\boldsymbol{v}})) \\ &= \boldsymbol{G}^{-1}\tilde{\boldsymbol{L}}\boldsymbol{Q}^T(\boldsymbol{I}_Q + \boldsymbol{Q}\boldsymbol{P}^T(\boldsymbol{I}_P - \boldsymbol{P}\delta\boldsymbol{G}\boldsymbol{G}^{-1}\tilde{\boldsymbol{L}}\boldsymbol{Q}^T\boldsymbol{Q}\boldsymbol{P}^T)^{-1}\boldsymbol{P}\delta\boldsymbol{G}\boldsymbol{G}^{-1}\tilde{\boldsymbol{L}}\boldsymbol{Q}^T)\boldsymbol{Q}(\delta\boldsymbol{h}' + \varrho(\delta\tilde{\boldsymbol{v}})) \\ &= \boldsymbol{G}^{-1}\tilde{\boldsymbol{L}}_Q(\boldsymbol{I}_Q + \boldsymbol{A}\boldsymbol{C})\boldsymbol{Q}(\delta\boldsymbol{h}' + \varrho(\delta\tilde{\boldsymbol{v}})) \end{aligned} \tag{S44}$$

where we defined

$$\tilde{\boldsymbol{L}}_Q := \tilde{\boldsymbol{L}}\boldsymbol{Q}^T \in \mathbb{R}^{N\times Q}, \tag{S45}$$
$$\boldsymbol{A} := \boldsymbol{Q}\boldsymbol{P}^T \in \mathbb{R}^{Q\times P}, \tag{S46}$$
$$\boldsymbol{B} := \boldsymbol{P}\delta\boldsymbol{G}\boldsymbol{G}^{-1}\tilde{\boldsymbol{L}}_Q \in \mathbb{R}^{P\times Q}, \tag{S47}$$
$$\boldsymbol{C} := (\boldsymbol{I}_P - \boldsymbol{B}\boldsymbol{A})^{-1}\boldsymbol{B} \in \mathbb{R}^{P\times Q}. \tag{S48}$$

This is a recurrence relation on $\delta\tilde{\boldsymbol{v}}$. To compute the perturbation response $\delta\boldsymbol{r}$ we simply note that, by Eq. S38,

$$\delta\boldsymbol{r} = \delta\boldsymbol{h}' + \varrho(\delta\tilde{\boldsymbol{v}}) + \boldsymbol{G}'\delta\tilde{\boldsymbol{v}} \tag{S49}$$
$$= f(\boldsymbol{v}^* + \delta\boldsymbol{h} + \delta\tilde{\boldsymbol{v}}) - \boldsymbol{r}, \tag{S50}$$

where we defined

$$\boldsymbol{G}' := \mathrm{diag}(f'(\boldsymbol{v}^* + \delta\boldsymbol{h})). \tag{S51}$$

Thus Eq. S44 can alternatively be written as a pair of recurrence relations on $\delta\tilde{\boldsymbol{v}}, \delta\boldsymbol{r}$ as

$$\delta\tilde{\boldsymbol{v}} = \boldsymbol{G}^{-1}\tilde{\boldsymbol{L}}_Q(\boldsymbol{I}_Q + \boldsymbol{A}\boldsymbol{C})\boldsymbol{Q}(\delta\boldsymbol{r} - \boldsymbol{G}'\delta\tilde{\boldsymbol{v}}), \tag{S52}$$
$$\delta\boldsymbol{r} = f(\boldsymbol{v}^* + \delta\boldsymbol{h} + \delta\tilde{\boldsymbol{v}}) - \boldsymbol{r}. \tag{S53}$$

Finally, instead of computing $\tilde{\boldsymbol{L}}_Q$ using its definition, which is inefficient as it requires computing a $N \times N$ matrix inverse, we compute its elements directly using the analytical solution for the linear perturbation response we previously derived[1]. Specifically, we compute the elements of $\tilde{\boldsymbol{L}}_Q$ by

$$[\tilde{\boldsymbol{L}}_Q]_{ij} \approx \frac{\pi}{p_{\alpha_j}\rho}\tilde{L}_{\alpha_i\alpha_j}(\mu_i, \mu_j, \boldsymbol{x}_i - \boldsymbol{x}_j, \theta_i - \theta_j) \tag{S54}$$

where $\alpha_i, \mu_i, \boldsymbol{x}_i, \theta_i$ are respectively the cell type, feature selectivity, spatial location, and feature tuning preference of neuron $i$, $\rho$ is the neuronal density, $p_\alpha$ is the fraction of neurons with cell type $\alpha$, and $\tilde{L}_{\alpha\beta}$ is given by[1]

$$\tilde{L}_{\alpha\beta}(\mu, \nu, \boldsymbol{x} - \boldsymbol{y}, \theta - \phi) := \frac{1}{\pi}\left(\tilde{L}_{0\alpha\beta}(\boldsymbol{x} - \boldsymbol{y}) + 2\tilde{L}_{1\alpha\beta}(\boldsymbol{x} - \boldsymbol{y})q(\mu)q(\nu)\cos(2(\theta - \phi))\right), \tag{S55}$$

where

$$\tilde{L}_{n\alpha\beta}(\boldsymbol{x}) := \frac{1}{2\pi}\sum_{\gamma=0}^{N_c^2-1} l_{n\alpha\beta\gamma}K_0\left(\sqrt{\lambda_{n\gamma}}\|\boldsymbol{x}\|\right), \tag{S56}$$
$$l_{n\alpha\beta\gamma} := [\tilde{\boldsymbol{U}}_n\boldsymbol{Q}_n]_{\alpha\gamma}[\boldsymbol{Q}_n^{-1}\tilde{\boldsymbol{V}}_n]_{\gamma\beta}, \tag{S57}$$

$N_c$ is the number of cell types, $\tilde{\boldsymbol{U}}_n \in \mathbb{R}^{N_c \times N_c^2}$, $\tilde{\boldsymbol{V}}_n \in \mathbb{R}^{N_c^2 \times N_c}$ are matrices defined by

$$\tilde{U}_{n\alpha\gamma} := \sum_{\beta=0}^{N_c-1} A_{n\alpha\beta}\delta_{N_c\alpha+\beta,\gamma}, \quad \tilde{V}_{n\gamma\beta} := \sum_{\alpha=0}^{N_c-1} \delta_{N_c\alpha+\beta,\gamma}$$
$$A_{0\alpha\beta} := g_\alpha w_{\alpha\beta}\sigma_{\alpha\beta}^{-2}, \quad A_{1\alpha\beta} := g_\alpha w_{\alpha\beta}\sigma_{\alpha\beta}^{-2}\kappa_{\alpha\beta}, \tag{S58}$$

(recall $g_\alpha$ is the neuronal gain of cell type $\alpha$), $\lambda_{n\gamma} \in \mathbb{C}$ and $\boldsymbol{Q}_n \in \mathbb{C}^{N_c^2 \times N_c^2}$ are defined such that $\boldsymbol{Q}_n \boldsymbol{\Lambda}_n \boldsymbol{Q}_n^{-1}$ is a diagonalization of $\boldsymbol{\Sigma}^{-1} - \tilde{\boldsymbol{V}}_n \boldsymbol{K}_n \tilde{\boldsymbol{U}}_n$, where $\boldsymbol{\Lambda}_n$ is the diagonal matrix of $\lambda_{n\gamma}$, $\boldsymbol{K}_n \in \mathbb{R}^{N_c \times N_c}$ is defined by

$$K_{0\alpha\beta} := \delta_{\alpha\beta}, \quad K_{1\alpha\beta} := \delta_{\alpha\beta}\mathbb{E}_\mu[q^2(\mu)] = \frac{2}{3}\delta_{\alpha\beta}, \tag{S59}$$

(see Methods for the last equality), and $\boldsymbol{\Sigma} \in \mathbb{R}^{N_c^2 \times N_c^2}$ is defined by

$$\Sigma_{\gamma\gamma'} := \delta_{\gamma\gamma'} \sum_{\alpha,\beta=0}^{N_c-1} \sigma_{\alpha\beta}^2 \delta_{N_c\alpha+\beta,\gamma}. \tag{S60}$$

Importantly, we note that $\tilde{\boldsymbol{L}}_Q, \boldsymbol{A}, \boldsymbol{C}, \boldsymbol{Q}$ all have $NQ$ elements or less, while $\boldsymbol{G}, \boldsymbol{G}'$ can be stored and multiplied as length-$N$ vectors since they are diagonal matrices. Thus, the space complexity of our approach is $\mathcal{O}(NQ)$ instead of the naive $\mathcal{O}(N^2)$, as claimed in Methods.

#### 5.2 Loss function

Formally, the loss function used for fitting the models in Figures 2-5 is given by

$$L = \sqrt{\frac{\sum_{k=0}^{K-1} w_k \sum_{n=0}^{N_k-1} \sigma_{kn}^{-2}(x_{kn} - y_{kn})^2}{\sum_{k=0}^{K-1} w_k \sum_{n=0}^{N_k-1} \sigma_{kn}^{-2} x_{kn}^2}} \tag{S61}$$

where $K$ is the number of datasets to which the models are fitted, $N_k$ is the number of data points in dataset $k$, $w_k$ is the weight of dataset $k$ in the loss function, $(x_{kn}, \sigma_{kn})$ are the mean and standard error of data point $n$ of dataset $k$ respectively, and $y_{kn}$ is the model prediction of that data point. Note that the loss is normalized such that $L = 1$ if $y_{kn} = 0$ for all $k, n$. More explicitly, $(x_{kn}, \sigma_{kn})$ and $y_{kn}$ are given by

$$x_{kn} := \langle x_{knp} \rangle_{p \in \mathcal{P}_k^{\text{data}}} \tag{S62}$$
$$\sigma_{kn} := \sqrt{\frac{\text{Var}_{p \in \mathcal{P}_k^{\text{data}}}(x_{knp})}{|\mathcal{P}_k^{\text{data}}|}} \tag{S63}$$
$$y_{kn} := \langle y_{knp} \rangle_{p \in \mathcal{P}_k^{\text{model}}} \tag{S64}$$

where $\mathcal{P}_k^X$ ($X \in \{\text{data}, \text{model}\}$) is a set of ensemble perturbations (possibly aggregated across different mice (when $X = \text{data}$) or different random instantiations of the model (when $X = \text{model}$)), and

$$x_{knp} := f_{k,n}(\delta \boldsymbol{r}_p^{\text{data}}) \tag{S65}$$
$$y_{knp} := f_{k,n}(\delta \boldsymbol{r}_p^{\text{model}}) \tag{S66}$$

for some scalar function $f_{k,n}$ of the perturbation response vector.

$K = 6$ for the models in Figures 2-3 and $K = 10$ for the models in Figures 4-5. For all the models, $w_k = 1$ for $k \in \{0, 1, 2, 3\}$, and $w_k = 0.25$ for $k \in \{4, 5\}$. $\mathcal{P}_0^X, \mathcal{P}_1^X, \mathcal{P}_2^X, \mathcal{P}_3^X$ are sets of diffuse untuned, compact untuned, diffuse cotuned, and compact cotuned ensembles respectively. More details regarding $\mathcal{P}_k^{\text{model}}$ can be found in Methods. Furthermore, $\mathcal{P}_4^X = \mathcal{P}_2^X$ and $\mathcal{P}_5^X = \mathcal{P}_3^X$. The functions $f_{k,n}$ are defined by

$$f_{k,n}(\delta \boldsymbol{r}) := \langle \delta r_i \rangle_{\{i|\alpha_i=\text{Pyr},d_i\in[15(n+1),15(n+2))\}} \tag{S67}$$

for all $(k, n) \in \{0, 1, 2, 3\} \times \{0, \cdots, 18\}$, and

$$f_{k,3s+t}(\delta \boldsymbol{r}) := \langle \delta r_i \rangle_{\{i|\alpha_i=\text{Pyr},d_i\in[15(s+1),15(s+2)),\Delta\theta_i\in[45t-22.5,45t+22.5),\mu_i\in[0.25,1)\}} \tag{S68}$$

for all $(k, s, t) \in \{4, 5\} \times \{0, \cdots, 8\} \times \{0, 1, 2\}$, where $\alpha_i$ is the cell type of neuron $i$, $d_i$ is the lateral distance of neuron $i$ from the nearest perturbed cell in µm, $\Delta\theta_i$ is the difference in preferred orientation of neuron $i$ from the perturbed ensemble, and $\mu_i$ is the orientation selectivity of neuron $i$.

For all $k \in \{6, 7, 8, 9\}$, $(x_{kn}, \sigma_{kn})$ are synthetic data points for regularizing the fitted models. Specifically, we set $x_{kn} = 0$ and $\sigma_{kn} = 0.01$ (in units of ΔF/F), where the value of $\sigma_{kn}$ is approximately the average standard error of all the data points in Figures S2C-D. $w_k = 3$ for all $k \in \{6, 7, 8, 9\}$, $\mathcal{P}_6^X, \mathcal{P}_7^X, \mathcal{P}_8^X, \mathcal{P}_9^X$ are equal to $\mathcal{P}_0^X, \mathcal{P}_1^X, \mathcal{P}_2^X, \mathcal{P}_3^X$ respectively, and $f_{k,n}$ is defined by

$$f_{k,3s+t}(\delta \boldsymbol{r}) := \begin{cases} \langle \delta r_i \rangle_{\{i|\alpha_i=\text{Pyr},d_i\in[250,\infty),\Delta\theta_i\in[45t-22.5,45t+22.5)\}}, & s = 0 \\ \langle \delta r_i \rangle_{\{i|\alpha_i=\text{PV},d_i\in[325,\infty),\Delta\theta_i\in[45t-22.5,45t+22.5)\}}, & s = 1 \\ \langle \delta r_i \rangle_{\{i|\alpha_i=\text{SST},d_i\in[325,\infty),\Delta\theta_i\in[45t-22.5,45t+22.5)\}}, & s = 2 \\ \langle \delta r_i \rangle_{\{i|\alpha_i=\text{VIP},d_i\in[325,\infty),\Delta\theta_i\in[45t-22.5,45t+22.5)\}}, & s = 3 \end{cases} \tag{S69}$$

for all $(k, s, t) \in \{6, 7, 8, 9\} \times \{0, 1, 2, 3\} \times \{0, 1, 2\}$. This regularization ensures that the responses of neurons far away from the perturbed ensemble are weak.

#### 5.3 Additional details about model constraints

Here we provide some additional details about the constraints imposed on the models during fitting. For constraint 17, the function $\tilde{L}_{\alpha\beta}(\boldsymbol{x})$ is given by[1]

$$\tilde{L}_{\alpha\beta}(\boldsymbol{x}) = \frac{1}{2\pi} \sum_{\gamma=0}^{N_c-1} [\boldsymbol{G}\boldsymbol{W}\boldsymbol{\Sigma}^{-1}\boldsymbol{Q}]_{\alpha\gamma} [\boldsymbol{Q}^{-1}]_{\gamma\beta} K_0 \left( \sqrt{\lambda_\gamma} \|\boldsymbol{x}\| \right) \tag{S70}$$

where $N_c$ is the number of cell types, $\boldsymbol{W}$ is the $N_c \times N_c$ matrix with elements $w_{\alpha\beta}$, $\boldsymbol{\Sigma}$ is the $N_c \times N_c$ matrix with elements $\sigma_{0\beta}^2$, and $\lambda_\gamma \in \mathbb{C}$, $\boldsymbol{Q} \in \mathbb{C}^{N_c \times N_c}$ are defined such that $\boldsymbol{Q}\boldsymbol{\Lambda}\boldsymbol{Q}^{-1}$ is a diagonalization of $(\boldsymbol{I} - \boldsymbol{G}\boldsymbol{W})\boldsymbol{\Sigma}^{-1}$, where $\boldsymbol{\Lambda}$ is the diagonal matrix of $\lambda_\gamma$.

Linear stability (constraint 8) is constrained in the same way as the models in Chau et al.[1]. We constrain that

$$\max_{k\in\mathcal{K}} \max_{n\in\{0,1\}} \max_{\lambda\in\sigma(\boldsymbol{J}_n(k))} \text{Re}(\lambda) < 0 \tag{S71}$$

where $\sigma(\boldsymbol{J}_n(k))$ denotes the set of eigenvalues of the $N_c \times N_c$ matrix $\boldsymbol{J}_n(k)$, $\mathcal{K}$ is the set of 1000 uniformly spaced points in the interval $\left[0, \frac{100}{\bar{\sigma}}\right]$ where $\bar{\sigma} := \sqrt{N_c^{-2} \sum_{\alpha\beta} \sigma_{\alpha\beta}^2}$, and the $(\alpha, \beta)$-th element of the matrix $\boldsymbol{J}_n(k)$ is given by

$$J_{n\alpha\beta}(k) := \tau_\alpha^{-1} \left( (1 + \sigma_{\alpha\beta}^2 k^2)^{-1} K_{n\alpha\alpha} A_{n\alpha\beta} \sigma_{\alpha\beta}^2 - \delta_{\alpha\beta} \right) \tag{S72}$$

where $K_{n\alpha\alpha}, A_{n\alpha\beta}$ are defined as in Eq. S58, S59.

### 6 Supplemental figures

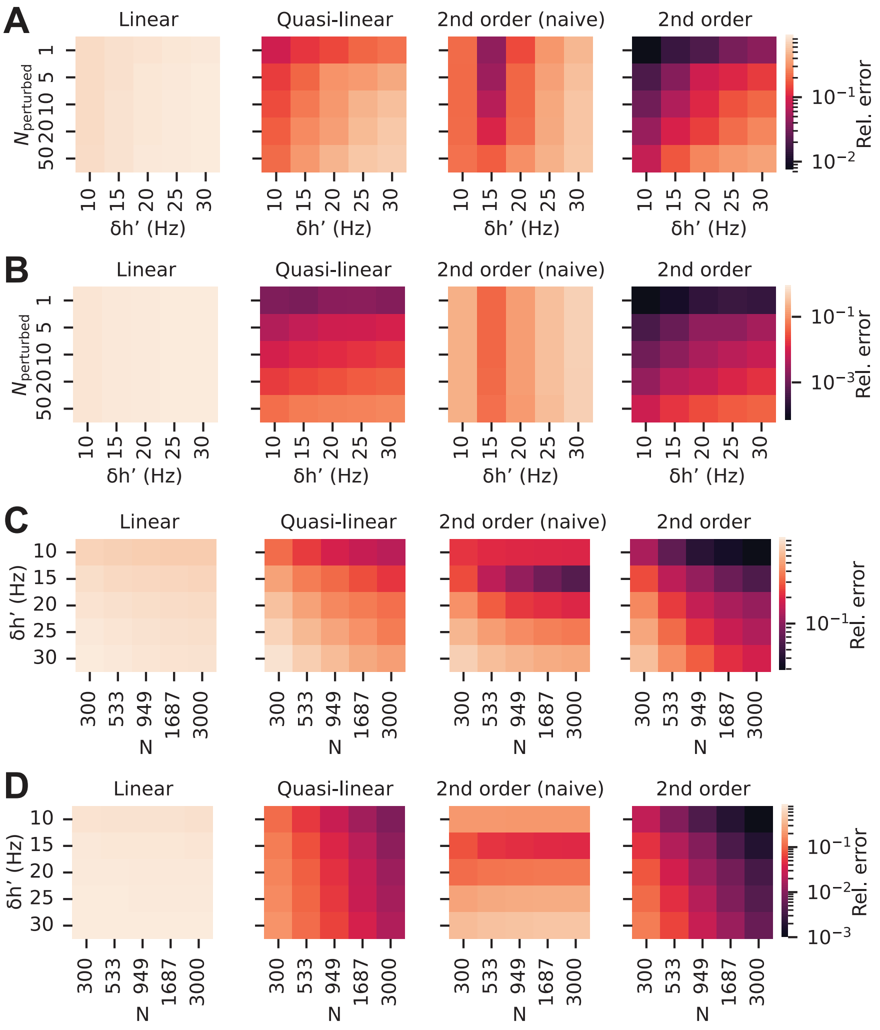


**Figure S1. Nonlinear theory of perturbation responses**

(A) Same as Figure 1C, but the relative errors of the linear approximation and the naive 2nd order approximation are also shown. Except for the case of $\delta h' = 15\,\mathrm{Hz}$, the relative error of the 2nd order approximation is substantially lower than that of the naive 2nd order approximation.
(B) Same as A, but the relative errors of the perturbed cells instead of the unperturbed cells are shown.
(C-D) Same as A-B, but the number of neurons in the random network instead of the number of perturbed cells is varied. The number of perturbed cells is 10.

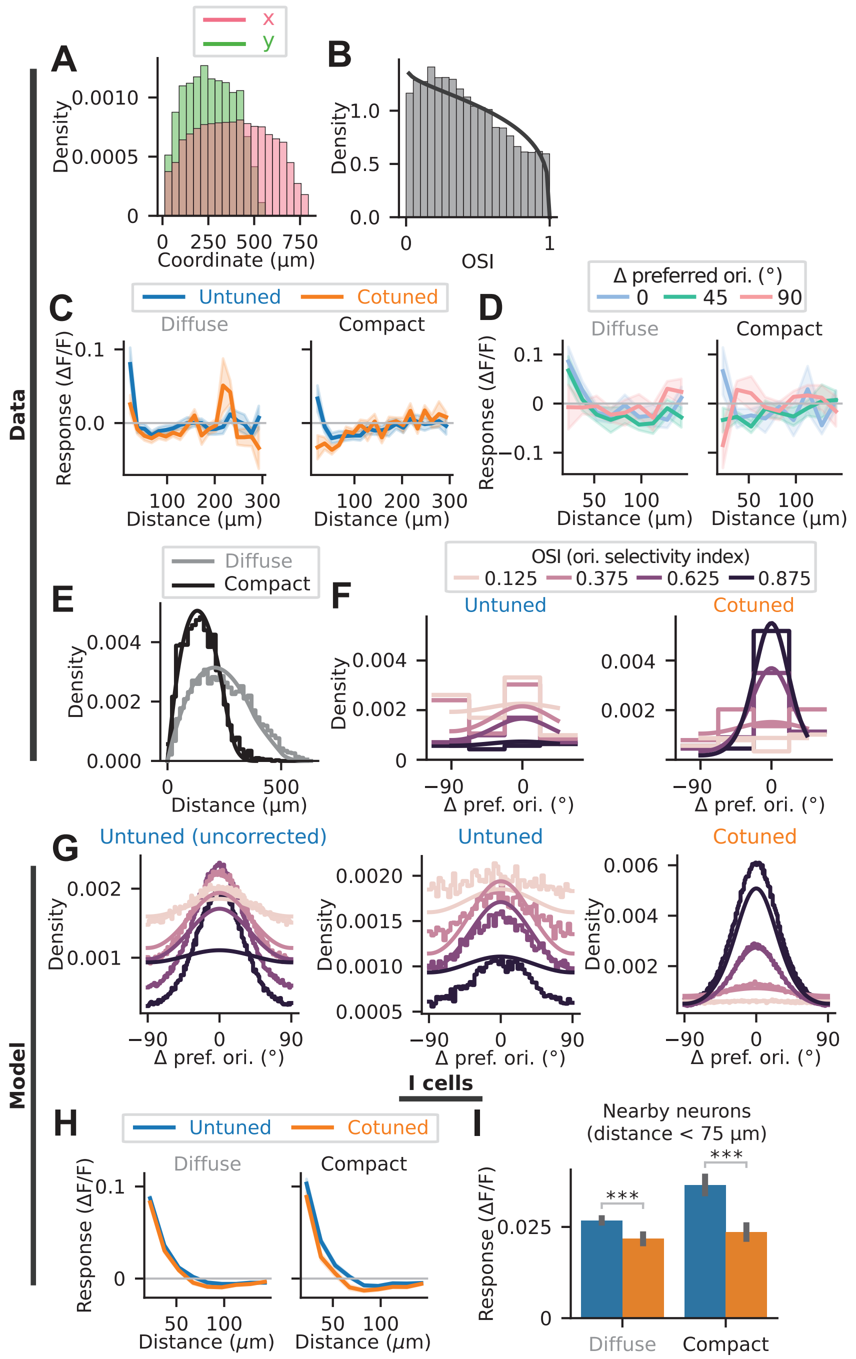


**Figure S2. Ensemble perturbations in mouse V1 L2/3 recruit nonlinear responses**

(A) Distribution of the spatial coordinates of the recorded cells in Oldenburg et al.[6]. Since most cells have an x coordinate between 0 and 750 µm and a y coordinate between 0 and 500 µm, we only analyze the responses of cells within a 750 µm × 500 µm spatial window in our models.
(B) Distribution of orientation selectivity index (OSI) of the recorded cells in Oldenburg et al.[6]. Solid line is the probability density function of the fitted Beta distribution $B(a_0, b_0)$, where $a_0 = 0.98$, $b_0 = 1.28$.
(C-D) Data points to which all models are fitted. C: Same as Figure 2D, but data points up to 300 µm are included. D: Perturbation response as a function of lateral cortical distance and Δ preferred orientation

for diffuse cotuned and compact cotuned ensembles in Oldenburg et al.[6]. Only the responses of visually responsive neurons with an OSI $> 0.25$ are included. Lines and shaded regions represent mean and standard error across ensembles.
(E) Distributions of pairwise distances between perturbed cells in diffuse and compact ensembles in Oldenburg et al.[6]. Smooth solid lines are the probability density functions of pairwise distances between points uniformly sampled from a $l_x \times l_y$ rectangle ($l_x = 504.9\,\mu\text{m}$, $l_y = 379.5\,\mu\text{m}$ for diffuse ensembles and $l_x = 292.4\,\mu\text{m}$, $l_y = 258.3\,\mu\text{m}$ for compact ensembles). $l_x$ and $l_y$ are obtained as the range between the $q$-th and $(1-q)$-th quantiles of the x and y spatial coordinates of the perturbed cells, where $q = 0.05$ for $l_x$ and $q = 0.1$ for $l_y$.
(F) Joint distributions of OSI and $\Delta$ preferred orientation (difference between the preferred orientation of the perturbed cells and the preferred orientation of the perturbed ensemble) of neurons in untuned and cotuned ensembles in Oldenburg et al.[6]. The joint distribution is visualized as histograms of $\Delta$ preferred orientation of perturbed cells in different OSI bins with bin width of 0.25. Smooth solid lines are the probability density functions of the fitted joint distribution, $f(\theta, \mu) := f_{\text{VM}}(\theta; 0, \exp(k_0 + k_1\mu + k_2\mu^2)) f_{\text{Beta}}(\mu; a_0 + a_1 - 1, b_0 + b_1 - 1)$, where $f_{\text{VM}}$ is the probability density function of the von Mises distribution, $f_{\text{Beta}}$ is the probability density function of the Beta distribution, $k_0 = -3.7, k_1 = 10.0, k_2 = -9.8, a_1 = 0.95, b_1 = 1.22$ for untuned ensembles, and $k_0 = -7.3, k_1 = 19.0, k_2 = -12.0, a_1 = 1.32, b_1 = 0.66$ for cotuned ensembles.
(G) Similar to F, but the histograms of $\Delta$ preferred orientation of perturbed cells in the model instead of the data are plotted. The OSI and $\Delta$ preferred orientation of perturbed cells in the model are effectively sampled from the fitted joint distribution (smooth solid lines), but our naive sampling scheme introduces a bias towards more cotuned ensembles (see Methods). This bias is particularly significant for untuned ensembles (left panel), but not so much for cotuned ensembles (right panel). By discarding untuned ensembles whose ensemble orientation selectivity is greater than 0.07, we approximately correct this bias (middle panel).
(H-I) Similar to Figure 2I-J, but the perturbation responses of inhibitory neurons in the model are shown. Notice that inhibitory cells in the model also exhibit cotuned-ensemble suppression ($^{***}p < 0.001$, one-sided exact paired permutation $t$-tests).

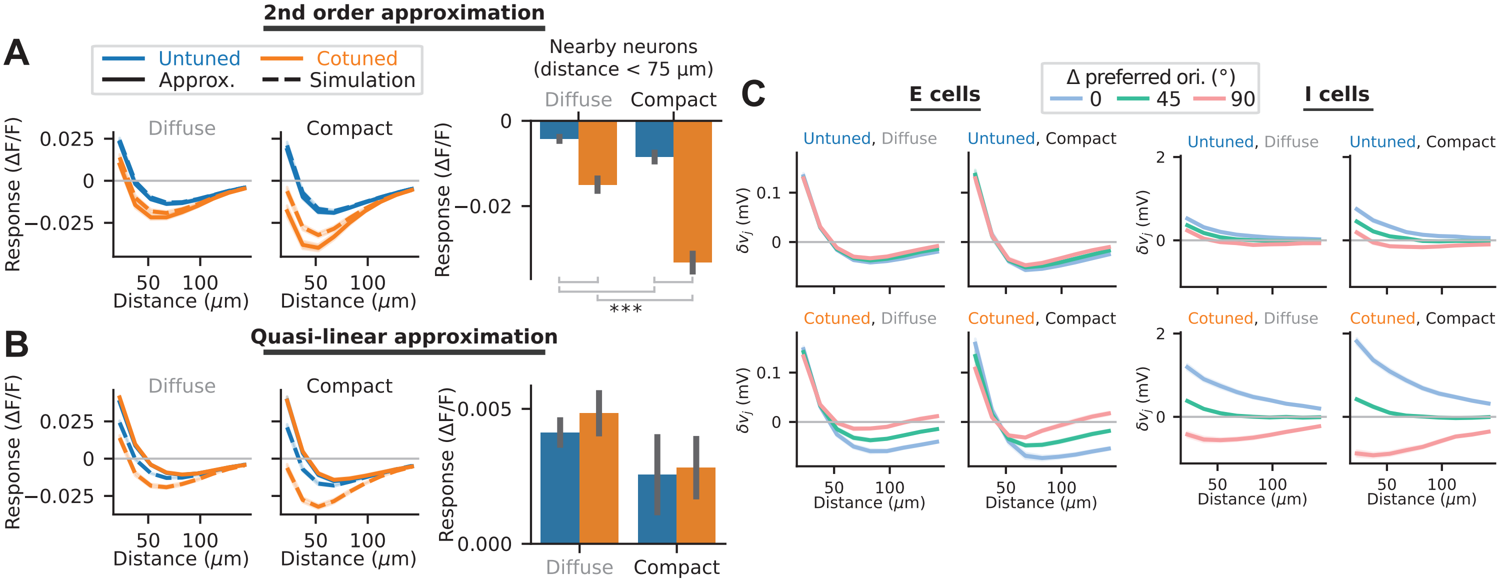


**Figure S3. A nonlinear inhibition pathway underlying cotuned ensemble perturbations**

(A) Left: Identical to Figure 3A, reproduced here for comparison. Right: Similar to Figure 2J, but the perturbation response is computed using the 2nd order approximation instead of numerical simulations. The 2nd order approximation captures the cotuned-ensemble suppression effect (***$p < 0.001$, one-sided exact paired permutation $t$-tests).

(B) Similar to A, but the quasi-linear approximation instead of the 2nd order approximation is used. Unlike the 2nd order approximation, the quasi-linear approximation cannot explain cotuned-ensemble suppression ($p > 0.05$, one-sided exact paired permutation $t$-tests).

(C) First-order change in recurrent input caused by ensemble perturbations in the best-fit nonlinear model, $\delta\boldsymbol{v}$, as a function of lateral cortical distance and $\Delta$ preferred orientation. $\delta v_j$ is much smaller for E cells (left) than for I cells (right), explaining why the (2,2,E) term of the 2nd order approximation, $\sum_{j\in\mathrm{E}} L_{ij}H_{jj}\delta v_j^2$, is negligible compared to the (2,2,I) term, $\sum_{j\in\mathrm{I}} L_{ij}H_{jj}\delta v_j^2$. Moreover, because the (2,1) term of the 2nd order approximation, $\boldsymbol{L}\delta\boldsymbol{G}\delta\boldsymbol{v}$, can be written as the sum $\sum_{j\in\mathcal{P}} L_{ij}\delta G_{jj}\delta v_j$, where $\mathcal{P}$ is the set of 10 E cells in the perturbed ensemble (since $\delta G_{jj} = 0$ for all $j \notin \mathcal{P}$), it also explains why the (2,1) term is negligible.

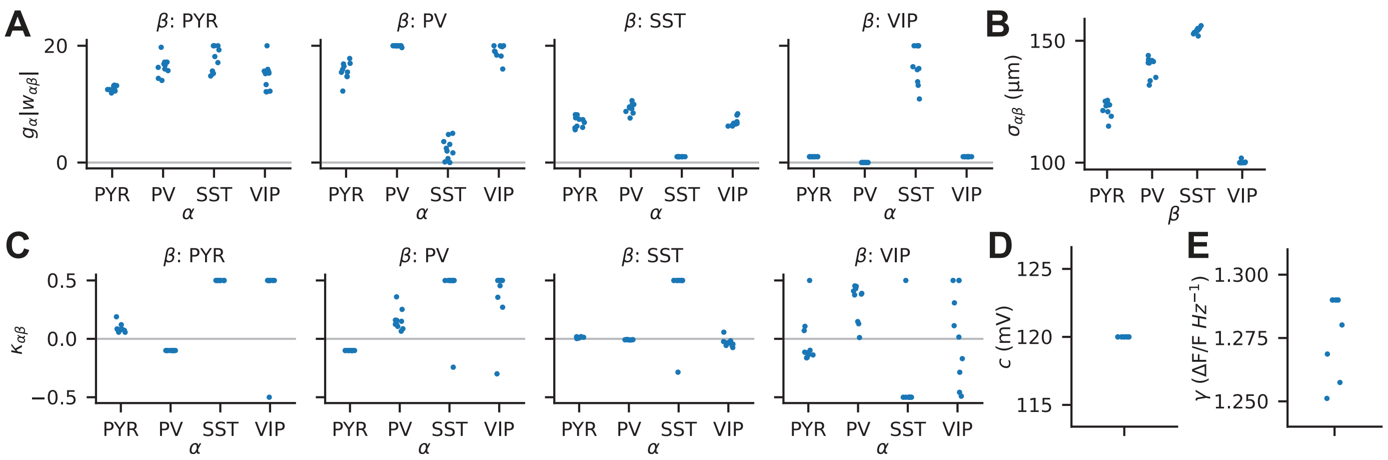


**Figure S4. SST as the predicted cell type mediating the nonlinear inhibition pathway**

(A-E) Model parameters of the top 10 best-fit models (one dot for each model). A: Linearized connection strength $g_\alpha|w_{\alpha\beta}|$, where $g_\alpha$ is the gain of cell type $\alpha$, and $g_\alpha|w_{\alpha\beta}|$ is constrained to be between 0 and 20. B: Connection widths $\sigma_{\alpha\beta}$, which are constrained to be independent of $\alpha$, such that there are only 4 instead of 16 parameters. $\sigma_{\alpha\beta}$ is also constrained to be between 100 µm and 175 µm. C: Connectivity tuning-dependence parameters $\kappa_{\alpha\beta}$, which are constrained to be between $-0.5$ and 0.5. Connections from cell type $\beta$ to cell type $\alpha$ are maximally like-to-like (neurons with similar tuning are more strongly connected) when $\kappa_{\alpha\beta} = 0.5$, and maximally like-to-unlike when $\kappa_{\alpha\beta} = -0.5$. D: Perturbation strength $c$, which is constrained to be between 80 mV and 120 mV (roughly corresponding to increases in perturbed cell firing rates by 8 Hz and 12 Hz respectively, since the transfer function of pyramidal cells is $f(v) = k[v]_+$, where $k = 0.1\,\mathrm{Hz\,mV^{-1}}$). E: Conversion factor from firing rate to ΔF/F, $\gamma$, constrained to be between 0 and 1.29.

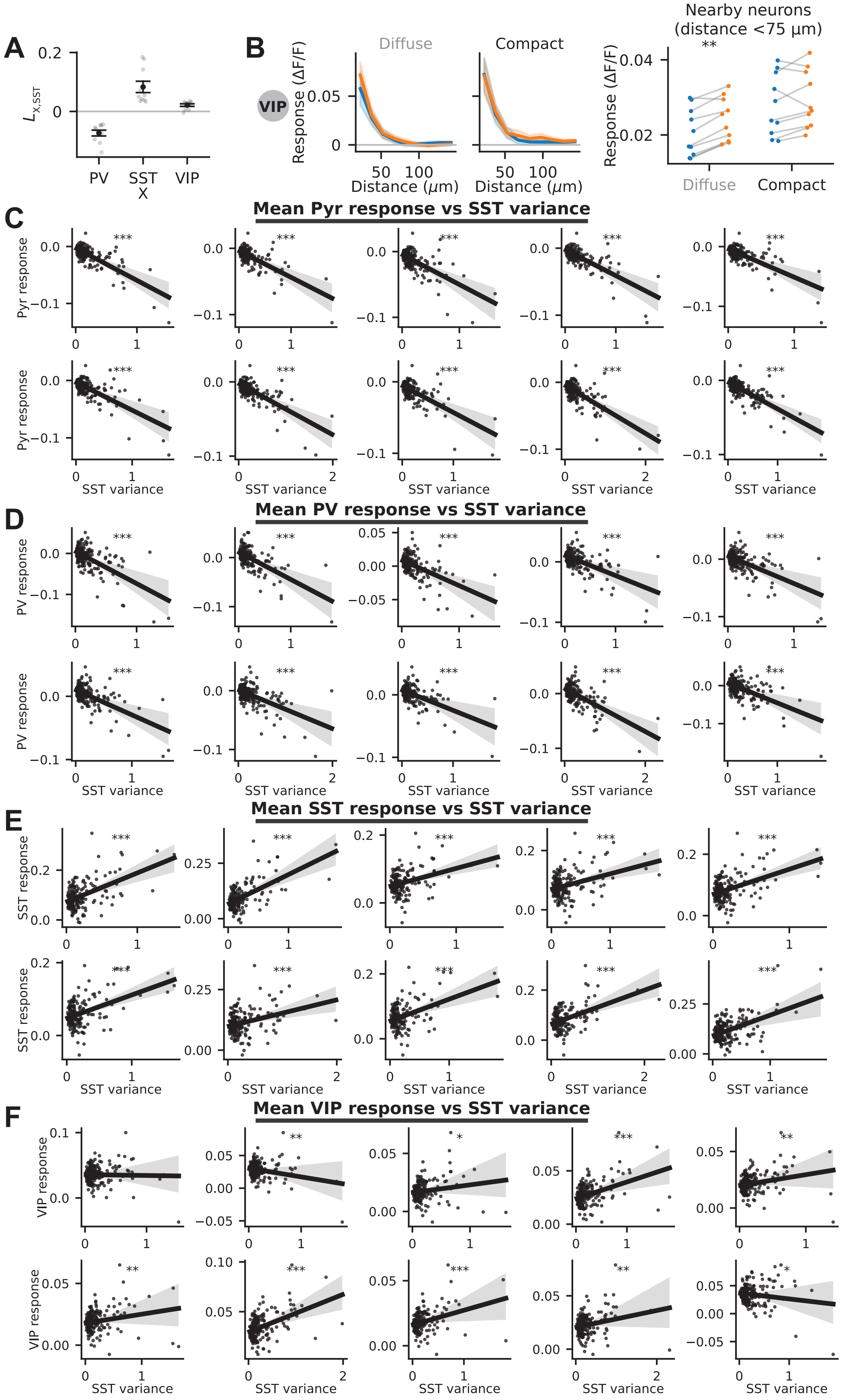
A
B
VIP
Diffuse
Compact
Nearby neurons (distance <75 µm)
Response (ΔF/F)
Distance (µm)
PV
SST
X
VIP
C
Mean Pyr response vs SST variance
Pyr response
SST variance
D
Mean PV response vs SST variance
PV response
SST variance
E
Mean SST response vs SST variance
SST response
SST variance
F
Mean VIP response vs SST variance
VIP response
SST variance

**Figure S5. Theory predicts cotuned-ensemble suppression of PV but facilitation of SST**

(A) Similar to Figure 5B, but including X = VIP.
(B) Similar to Figure 5C, but for VIP neurons ($^{**}p = 0.004$, two-sided exact Wilcoxon signed-rank tests).
(C-F) Similar to Figure 5E, but the correlations between SST variances and the mean responses of the 4 cell types are shown for all top 10 best-fit models (one subplot per model) ($^{*}p < 0.05$, $^{**}p < 0.01$, $^{***}p < 0.001$, two-sided permutation tests of Pearson correlation coefficient).

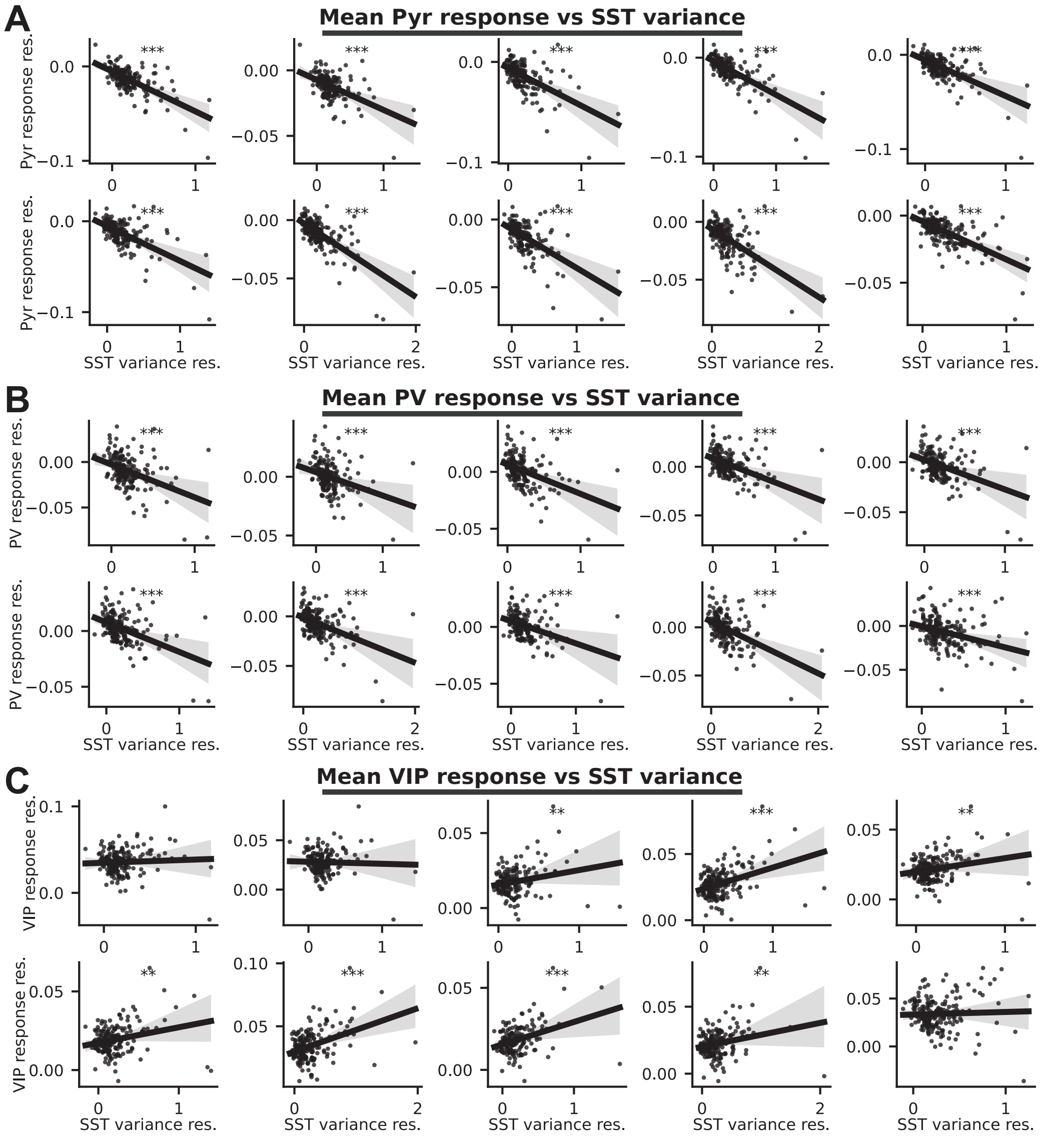


**Figure S6. Theory predicts cotuned-ensemble suppression of PV but facilitation of SST (cont.)**

(A-C) Correlation of the mean responses of nearby Pyr, PV, and VIP neurons ($< 75\,\mu m$) with the variances of nearby SST activity ($< 75\,\mu m$) after regressing out the mean responses of nearby SST neurons ($< 75\,\mu m$), for all top 10 best-fit models (one subplot per model). The regression on mean nearby SST responses is applied to both the x and y axis variables, and the means of both variables are added back to the residuals to keep the scale of each axis comparable to Figure S5C-F. Each dot corresponds to an ensemble perturbation. Lines represent linear regression fits, and shaded regions represent the 95% confidence intervals obtained by bootstrapping with 9999 resamples ($^{*}p < 0.05$, $^{**}p < 0.01$, $^{***}p < 0.001$, two-sided permutation tests of Pearson correlation coefficient).
